# Transcription factor NFYA directs male meiotic entry by facilitating accessible chromatin at meiotic promoters in mice

**DOI:** 10.1101/2025.05.09.653044

**Authors:** Martin Säflund, Masomeh Askari, Atiyeh Eghbali, Mukhtar Mohamed Abdi, Ann-Kristin Östlund Farrants, Tianxiong Yu, Deniz M. Özata

## Abstract

Meiotic prophase I characterized by homologous recombination and synapsis is an intricate step for spermatogenesis. This process entails extensive changes to chromatin and transcription. Prior to prophase I, accessible chromatin bound by paused Pol II at meiotic gene promoters is essential for their timely activation later during meiosis. However, the factor responsible for promoting accessible chromatin at meiotic gene promoters before entry into prophase I is unknown. Here, we discovered that NFYA expressed in pre-meiotic germ cells promotes accessible chromatin at meiotic gene promoters including those regulated by STRA8/MEISON axis. Concordantly, conditional germline deletion of *Nfya* in male mice blocks meiotic entry. Functionally, our spatial and single-cell ATAC-seq data revealed that loss of NFYA in pre-meiotic cells disrupts accessible chromatin at meiotic gene promoters. Our study identifies a pioneer role for NFYA in facilitating accessible chromatin at meiotic gene promoters before meiosis, thereby regulating the timely activation of meiotic genetic program.

## INTRODUCTION

Mammalian spermatogenesis is a stepwise developmental process to produce spermatozoa from spermatogonial stem cells (SSCs)^1,2^. This intricate and highly conserved developmental program involves three major steps. Mitotically dividing SSCs differentiate and enter meiosis^3^; upon meiotic entry, primary spermatocytes (SpIs) undergo two rounds of meiotic division resulting in haploid round spermatids (RSs)^1,2^; RSs then finally differentiate into spermatozoa, i.e., spermiogenesis^4^. Meiosis is a critical step during spermatogenesis. Upon entry into prophase I of meiosis, chromosomes are reorganized into proteinaceous structures, axial elements (AE), to facilitate homologous chromosome pairing and synapsis, as well as crossover^1,2,5^.

Both entry into meiosis and progression through prophase I entails extensive chromatin and transcriptional changes^6–9^. The switch from mitosis to meiosis is regulated by the transcription factors (TFs) STRA8 and MEIOSIN, which activate the meiotic genetic program in response to retinoic acid^10,11^. During this switch, Znhit1 replaces canonical histone H2A with H2A.Z^12^. At the pachytene stage of prophase I, the TFs A-MYB and TCFL5 initiate the transcriptional burst of thousands of meiotic genes, including genes producing pachytene piRNAs, as well as genes required during spermiogenesis^13–17^.

Strikingly, in pre-meiotic spermatogonia (SpG), the promoters of genes expressed during meiosis already have accessible chromatin^18^, and are occupied by paused RNA polymerase II (Pol II)^19^. Establishment of paused Pol II at gene promoters in spermatogonia is thought to be a key regulatory step that primes genes for timely activation later during meiosis. In fact, at pachytene stage of prophase I, A-MYB recruits testis-specific bromodomain protein, BRDT, to release paused Pol II into elongation^20^. Nevertheless, the factor that likely conducts the pioneering activity required to establish accessible chromatin, and thus facilitate the binding of Pol II at meiotic gene promoters before entry into prophase I, remains unknown.

Here, we demonstrate that prior to the initiation of meiosis, NFYA promotes accessible chromatin at the promoters of genes expressed during meiotic entry and progression. Germline-specific deletion of *Nfya* results in spermatogenic arrest at meiotic entry. Together, our findings identify NFYA as a pioneer factor regulating chromatin accessibility during spermatogenesis and provide new insights to the molecular mechanisms of meiotic initiation in mice.

## RESULTS

### Each major step of spermatogenesis retains a distinct gene expression profile

To define the expression profiles of genes across spermatogenesis, we used our published RNA sequencing (RNA-seq) data from FACS-purified germ cells with spike-in sequences^13,17,21^, and classified genes according to their absolute steady-state transcript abundance: i.e., molecules per FACS-purified germ cell (Figures 1A and S1A; Table S1A). Those genes, with transcript abundance ≥4-fold higher in spermatogonia (SpG) than in pachytene/diplotene spermatocytes (P/D), secondary spermatocytes (SpII), and round spermatids (RS), were classified as mitosis genes (1,181 genes). Whereas, the transcript abundance of meiosis-I genes in P/D was ≥4-fold higher than in other germ cells (3,062 genes). As RNAs required for spermiogenesis are first expressed during prophase I^22–25^, we classified the third category as spermiogenesis genes whose transcript abundance in P/D was ≥4-fold higher than in SpG and changed ≤2-fold in SpII and RS (2,345 genes). Exemplified genes confirmed the differential expression of previously described marker genes across spermatogenesis (Figures S1B–S1D). The mRNA abundance of *Sall4* and *Dmrt1* genes were highest in SpG consistent with their crucial function in the maintenance spermatogonia^26,27^ (Figure S1B). In human and mouse testis, ∼100 well-annotated genes produce long transcripts that are processed into pachytene piRNAs, i.e., pachytene piRNA genes^14,15^. The transcription of pachytene piRNA genes is initiated by the collaborative activity of A-MYB and TCFL5 during pachytene stage of male prophase I^14,16,17^. Consistently, the piRNA precursor transcript of *pi9* was highest in P/D cells (Figure S1C). Of note, the category for meiosis-I genes included 99 of the 100 mouse pachytene piRNA genes (Tables S1A and S1B). mRNA of *sycp2* gene, whose protein product is an essential component of the synaptonemal complex formed during prophase I of meiosis^28^, was higher in P/D compared to other germ cells (Figure S1C). *Crem* and *Spata6* genes are essential for spermiogenesis^29,30^. Consistently, they were first expressed in P/D cells and their RNA abundance increased in RS cells (Figure S1D).

**Figure 1.**
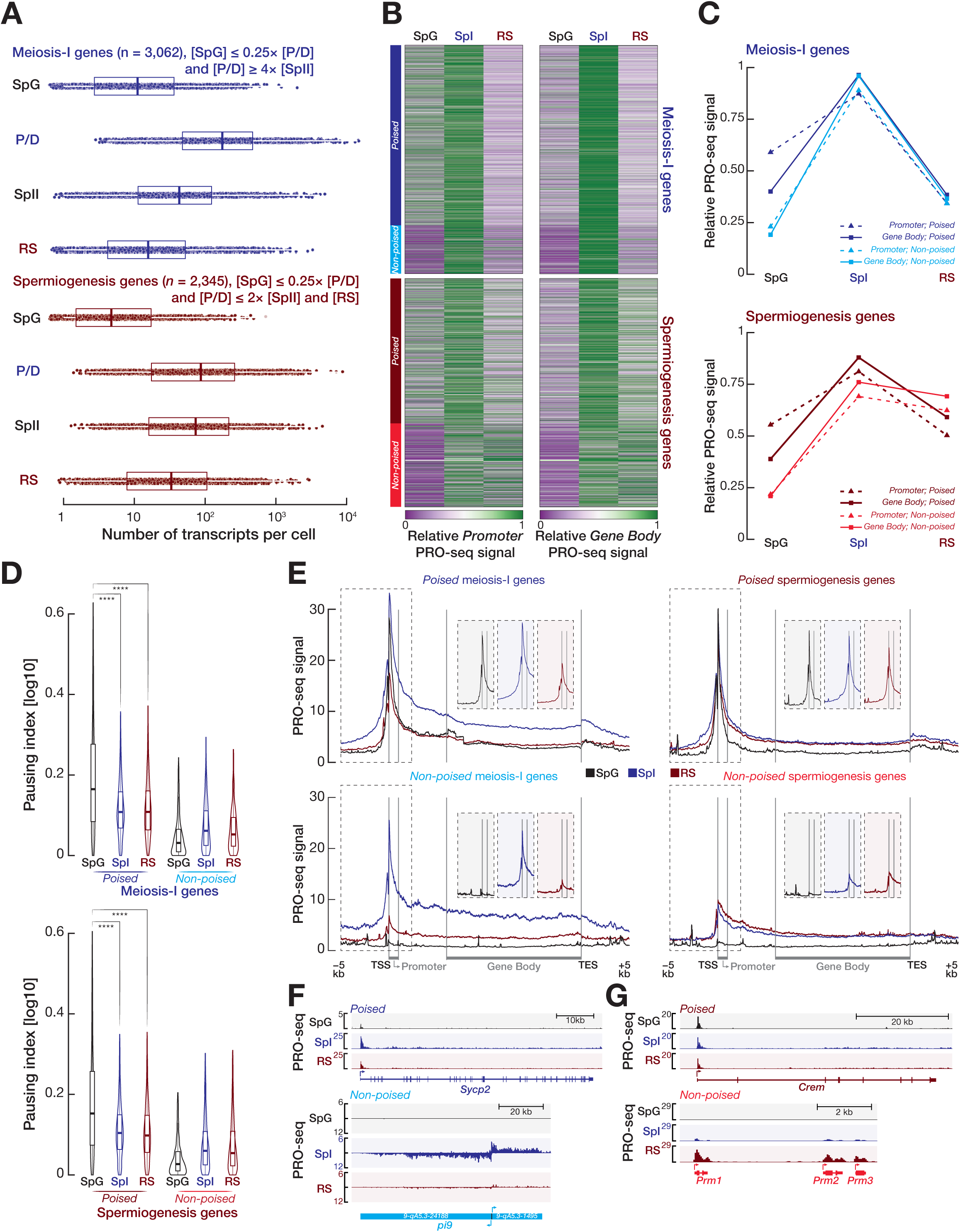
Many meiosis-I and spermiogenesis genes accumulate paused Pol II around their TSSs before meiotic entry. **(A)** Box plots show absolute steady-state transcript abundance of meiosis-I and spermiogenesis genes in spermatogonia (SpG), primary (SpI) and secondary (SpII) spermatocytes, and round spermatids (RS). Vertical lines represent median. Whiskers represent maximum and minimum values. Data further than 1.5 × interquartile range (IQR) represents outliers. **(B)** Heatmaps show the relative promoter PRO-seq signal (left; from transcription start site [TSS] to first 5% of gene) and relative gene body PRO-seq signal (right; from the first 30% of the gene to transcription end site [TES]) across SpG, SpI, and RS cells. **(C)** Line plots (top, meiosis-I genes; bottom, spermiogenesis genes) represent the average relative promoter (dashed line with triangle) and gene body (line with square) PRO-seq signals for poised and non-poised genes across SpG, SpI, and RS cells. **(D)** Violin plots (top, meiosis-I genes; bottom, spermiogenesis genes) of pausing index calculated from the ratio of promoter PRO-seq density to gene body PRO-seq density on poised and non-poised genes. \*\*\*\**p* < 0.0001; two-sided wilcoxon matched-pairs signed ranked sum test. Horizontal lines represent median. Whiskers show maximum and minimum values. IQR is represented by boxplots. Plotted pausing indexes are the average of three biological replicates from SpG, SpI, and RS cells. **(E)** Metagene plots (left two plots, meiosis-I genes; right two plots, spermiogenesis genes) of average PRO-seq signals of three biological replicates from SpG, SpI, and RS cells at annotated gene boundaries for poised and non-poised genes. Insets show PRO-seq density for SpG, SpI, and RS at the promoters (from TSS to first 5% of gene length). **(F, G)** Integrative Genomics Viewer (IGV) views for PRO-seq signals at gene boundaries of exemplified poised and non-poised meiosis-I genes **(F)** and spermiogenesis genes **(G)**. See also Figures S1 and S2.

We further performed Gene Ontology analysis (GO; ≥2-fold change; Fisher’s exact test, FDR < 0.01; Tables S1C–S1E) to characterize the biological function of genes in each category. We found that mitotic genes were enriched for mitotic DNA replication, housekeeping functions such as cell proliferation and morphogenesis, and macromolecule biosynthesis (Table S1C). GO for meiosis-I genes revealed enriched gene categories related to meiosis, including positive regulation of DNA repair, male meiotic nuclear division, and piRNA processing (Figure S1E; Table S1D). Notably, spermiogenesis genes were mainly enriched for GO related to spermatid differentiation, sperm function, and fertilization such as acrosome assembly, flagellated sperm movement, and sperm-egg recognition (Figure S1E; Table S1E).

### Promoter-proximal regions of genes expressed during meiosis accumulate paused Pol II in spermatogonia

Recent studies reported that spermatogonia and early prophase I cells, leptotene/zygotene, accumulate paused RNA polymerase II (Pol II) around the transcription start sites (TSSs) of genes expressed during meiosis. Pol II pausing at TSSs of such genes is required for the timely transcriptional activation during the pachynema stage of prophase I^19,20^. In fact, A-MYB, expressed in pachytene cells^14,31^, recruits BRDT to release paused Pol II into elongation^20^.

We sought to quantify paused and elongating Pol II within meiosis-I and spermiogenesis genes across spermatogenesis using publicly available <u>P</u>recision <u>R</u>un- <u>O</u>n sequencing (PRO-seq) data from purified spermatogonia (SpG), primary spermatocytes (SpI), and round spermatids (RS) cells^19^. Spike-in sequences in each sample enabled us to quantify absolute PRO-seq signal which is corrected for genome copy number.

Engaged Pol II peak around TSS is the characteristic of promoter-proximal pausing of Pol II^32^, wherein the promoters of many metazoan developmental genes exhibit paused Pol II that is later released into active elongation in response to secondary signals^33^. We thus first assessed whether the TSSs of meiosis-I and spermiogenesis genes retain significant Pol II peaks in SpG cells prior to their expression in SpI or RS cells. Using MACS3 (FDR < 0.01), we found that ∼79% of meiosis-I genes exhibited significant Pol II peaks within the ±2 kb of their TSSs in SpG cells (2,415 of 3,062 genes; median distance from the TSS to nearest Pol II peak = 158 bp) (Figure S2A; Table S1B). Likewise, the promoters of ∼63% of spermiogenesis genes retained Pol II peak in SpG cells (1,488 of 2,345 genes; median distance from the TSS to nearest Pol II peak = 166 bp) (Figure S2A; Table S1B). This suggests that Pol II is loaded on chromatin and remains paused at the promoters of the majority of both meiosis-I and spermiogenesis genes prior to their expression in SpI or RS cells (Figure S2A). Therefore, we named those genes with Pol II peak near their TSSs as poised genes, while genes without significant Pol II peak within the ±2 kb of their TSSs were named non-poised genes. We next measured the relative promoter and gene body PRO-seq signal for poised and non-poised genes across SpG, SpI and RS cells (Figures 1B and 1C). For both poised and non-poised meiosis-I genes, active RNA synthesis at gene bodies was highest in SpI cells, and declined ∼3-fold in RS cells (Figures 1B and 1C). Whereas, nascently transcribed RNA signal at gene bodies from poised and non-poised spermiogenesis genes remained almost unchanged between SpI and RS cells (decrease <2-fold; Figures 1B and 1C). Mitosis genes, by contrast, revealed a ∼3-fold higher promoter and gene body PRO-seq signal in SpG than in SpI and RS cells, suggesting that their transcription is switched off after entry into meiosis I (Figures S2B and S2C). Importantly however, although both poised and non-poised genes exhibited maximal gene body PRO-seq signal in SpI or RS cells compared with SpG cells, promoter PRO-seq signal of poised genes was >2-fold higher than that of non-poised genes in SpG cells (Figures 1B and 1C), consistent with the Pol II peak near the TSSs of poised genes in SpG cells (Figure S2A).

To quantitatively assess pause-release across each cell stage, we calculated the polymerase pausing index (i.e., the density of PRO-seq signal at promoter relative to the PRO-seq density at gene body) for poised and non-poised genes. Intriguingly, pausing index for poised genes in SpG cells was ∼2-fold higher than in SpI and RS cells (Two-sided wilcoxon matched-pairs signed ranked sum test; \*\*\*\**p* < 0.0001), whereas we did not observe a higher pausing index for non-poised genes in SpG cells when compared to SpI and RS cells (Figure 1D). Metagene plots corroborated computed pausing indexes for poised and non-poised genes: In SpG cells, poised genes displayed high Pol II signal at promoters, but not at gene bodies, whereas proximal-promoter PRO-seq signal for non-poised genes in SpG cells was at the background level (Figure 1E). For example, poised meiosis-I (*sycp2*) and spermiogenesis (*Crem,* required for spermatid development^29^) genes had high levels of promoter-proximal Pol II accumulation in SpG cells although active RNA production from their gene bodies ensued later in SpI or RS cells (Figures 1F and 1G). Non-poised meiosis-I (*pi9*, pachytene piRNA gene^14^) and spermiogenesis (*Prms*, encoding protamines essential for spermatozoa chromatin condensation^34^) genes, on the other hand, lacked accumulation of Pol II around their TSSs in SpG cells, while they are actively transcribed in SpI or RS cells (Figures 1F and 1G).

Finally, we sought to test whether A-MYB and BRDT occupancies around the TSSs of poised genes are higher than that of non-poised genes in pachytene spermatocytes using our previously published <u>C</u>utting <u>U</u>nder <u>T</u>arget and <u>R</u>elease <u>U</u>sing <u>N</u>uclease (CUT&RUN;^35^) for A-MYB^13,17^ and publicly available BRDT ChIP-seq data^36^. We found that in pachytene spermatocytes, the promoters of poised meiosis-I genes had higher A-MYB (median = 7.4 rpm) and BRDT (median = 4.5 rpm) occupancies compared to that of non-poised meiosis-I genes (median A-MYB occupancy = 3.2 rpm; median BRDT occupancy = 3.6 rpm) (Figure S2D). However, while A-MYB occupancy was higher at the promoters of poised spermiogenesis genes compared with that of non-poised genes, we did not observe higher BRDT occupancy at the promoters of poised spermiogenesis genes (Figure S2D). This suggests that for poised spermiogenesis genes, the release of Pol II into active elongation likely depends on other BET proteins or factors. Together, we conclude that in spermatogonia, the promoters of majority meiosis-I and spermiogenesis genes accumulate paused Pol II, thereby remaining poised for activation during later meiosis.

### Promoters of poised genes have accessible chromatin in spermatogonia

Pol II pausing at developmentally regulated genes is among the major regulatory steps allowing for the expression of such genes in a correct spatiotemporal sequence. Yet chromatin requires accessible state for Pol II to bind DNA (Reviewed in^37^). To test the idea that accumulation of paused Pol II at poised gene promoters is accompanied by accessible chromatin in spermatogonia, we purified populations of spermatogonia (SpG), leptonene/zygotene (L/Z), and pachytene/diplotene (P/D) cells from wild-type C57BL/6 mice with high purity using FACS (Figure S3) and performed <u>A</u>ssay for <u>T</u>ransposase-<u>A</u>ccessible <u>C</u>hromatin using sequencing (ATAC-seq). Intriguingly, in SpG cells, the chromatin around promoters of poised genes showed strong ATAC-seq signal which is sustained throughout prophase I, whereas the promoters of non-poised genes lacked ATAC-seq signal for accessible chromatin in this stage (Figures 2A and S4A). However, the promoters of non-poised genes displayed ATAC-seq signal in only P/D cells, consistent with their expression at this stage (Figures 2A and S4A). To ensure robust conclusion that poised meiosis-I and spermiogenesis genes have accessible chromatin around their promoters in SpG cells, we located significant ATAC-seq peaks genome-wide using MACS3 (FDR < 0.01; Table S2A) and computed the distance from the TSSs of poised and non-poised genes to nearest ATAC-seq peak in three biological replicates from each cell stage (Figure S4B; agreement between replicates; SpG Spearman’s ρ > 0.83, L/Z Spearman’s ρ > 0.92, P/D Spearman’s ρ > 0.95). Genes whose promoters exhibit ATAC-seq peaks within the ±2 kb of their TSSs in at least two replicates were considered as genes with an accessible promoter (Figure 2B). Strikingly, in SpG cells, >90% of poised genes had ATAC-seq signal near their TSSs (poised meiosis-I genes, median distance from TSS to nearest ATAC-seq peak = 186 bp; poised spermiogenesis genes, median distance = 182 bp), whereas, the TSSs of ≤40% of non-poised genes displayed ATAC-seq peak (non-poised meiosis-I genes, median distance = 6,456 bp; non-poised spermiogenesis median distance =10,233 bp) (Figure 2B; Table S2B). Exemplified poised and non-poised genes corroborated our finding: the promoters of poised genes exhibited significant ATAC-seq peak near their TSSs, whereas the promoters of non-poised genes lacked ATAC-seq signal (Figures 2C and 2D). Finally, we asked whether, in both SpG and SpI cells, chromatin accessibility at promoters predicted the engaged Pol II peak around the TSSs of genes we categorized. We found moderate to strong correlation between the accumulation of Pol II signal and chromatin accessibility at meiosis-I and spermiogenesis gene promoters (Figures 2E and 2F; Spearman’s ρ = 0.38–0.72). We conclude that accessible chromatin around the promoters of poised genes is established in pre-meiotic cells prior to the onset of their transcription during prophase I of meiosis.

**Figure 2.**
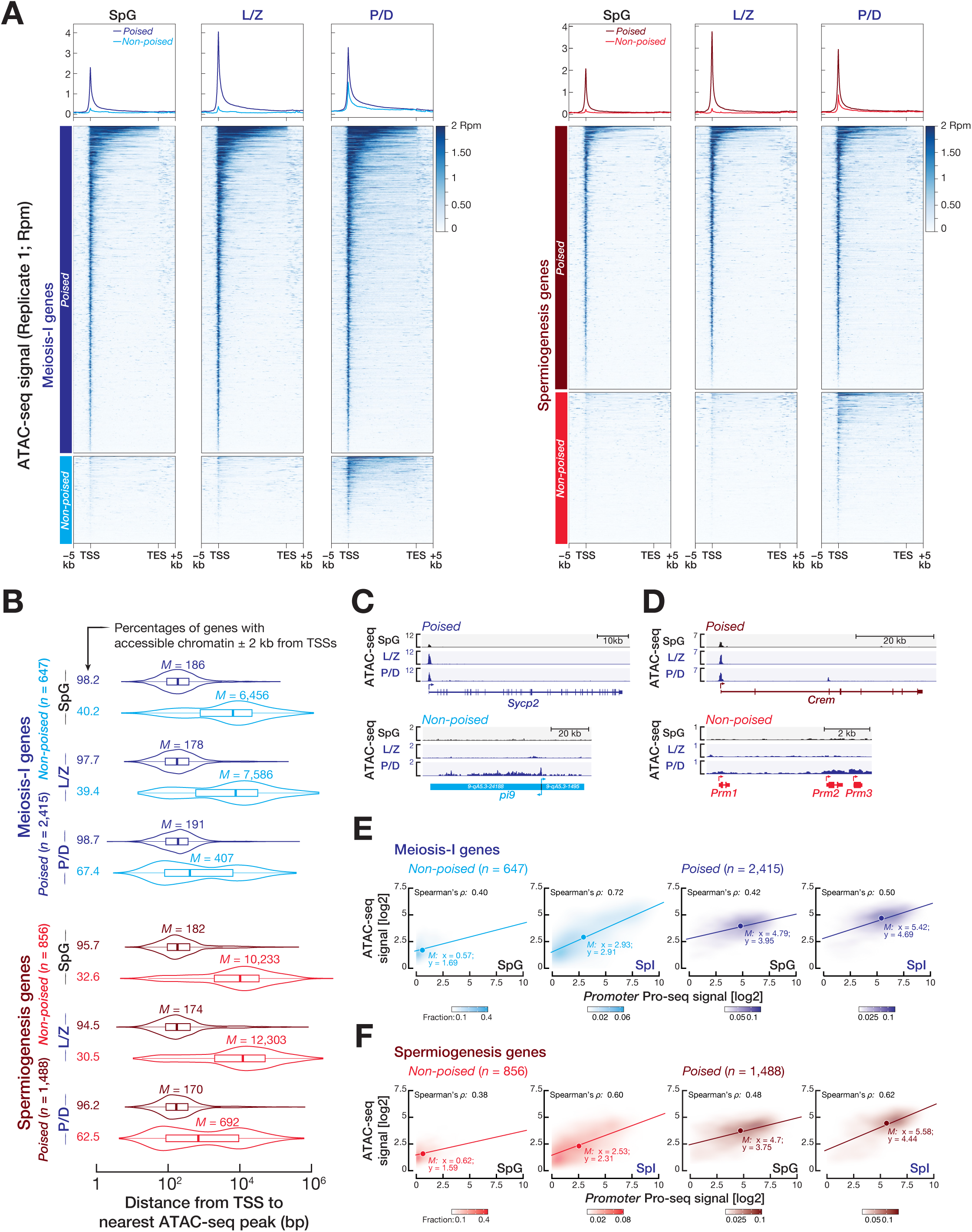
Chromatin is in a permissive state at the promoters of poised genes in spermatogonia. **(A)** Metagene plots (top) and heatmaps (bottom) show reads per million (Rpm)-normalized ATAC-seq signals in the –5 kb to +5 kb window flanking transcription start sites (TSSs) and transcription end sites (TESs) of poised and non-poised genes for spermatogonia (SpG), leptotene/zygotene (L/Z), and pachytene/diplotene (P/D) cells. **(B)** Distance (average of three biological replicates) from the TSS to nearest ATAC-seq peak for poised and non-poised meiosis-I and spermiogenesis genes in SpG, L/Z, and P/D cells. Genes were classified as genes with accessible chromatin at promoters if they had significant ATAC-seq peak ± 2 kb from their TSSs in at least two biological replicates of ATAC-seq experiment. Vertical lines represent median. Whiskers show maximum and minimum values. IQR is represented by boxplots. **(C, D)** Integrative Genomics Viewer (IGV) views for ATAC-seq signals at gene boundaries of exemplified poised and non-poised meiosis-I genes **(C)** and spermiogenesis genes **(D)**. **(E, F)** Scatter plots show the spearman’s correlation (*P*) between ATAC-seq density and PRO-seq density at the promoters of poised and non-poised meiosis-I genes **(E)** and spermiogenesis genes **(F)** for SpG and SpI cells. Each data point represents average of three biological replicates. See also Figures S3 and S4.

### Accessible poised gene promoters retain binding motif for NFYA that is highly expressed in pre-meiotic cells

An increasing number of TFs have been shown to both bind *cis*-regulatory regions in compact chromatin, and have the ability to promote chromatin opening—thereby facilitating the subsequent binding of additional factors, such as general TFs, Pol II, and tissue-specific TFs. Such factors are collectively called pioneer factors. The function of pioneer factors is critical for the timely expression of developmental genes (Reviewed in ^38^). We thus searched for enriched TF binding motifs under ATAC-seq peaks located around the promoters of poised genes in spermatogonia (HOMER; Benjamini-Hochberg multiple test correction, FDR < 0.05). NFYA binding motif, RRCCAATSRS, was significantly enriched as one of the top scoring motifs beneath the ATAC-seq peaks around the promoters of poised genes in spermatogonia (Figure 3A). The CCAAT pentanucleotide located near TSSs is bound by several TFs including NFYA, CTF (CCAAT Transcription Factor), and C/EBP (CCAAT Enhancer Binding Protein)^39^. Notably, NFYA maintains the chromatin accessibility around the TSSs in embryonic stem cells^40^.

**Figure 3.**
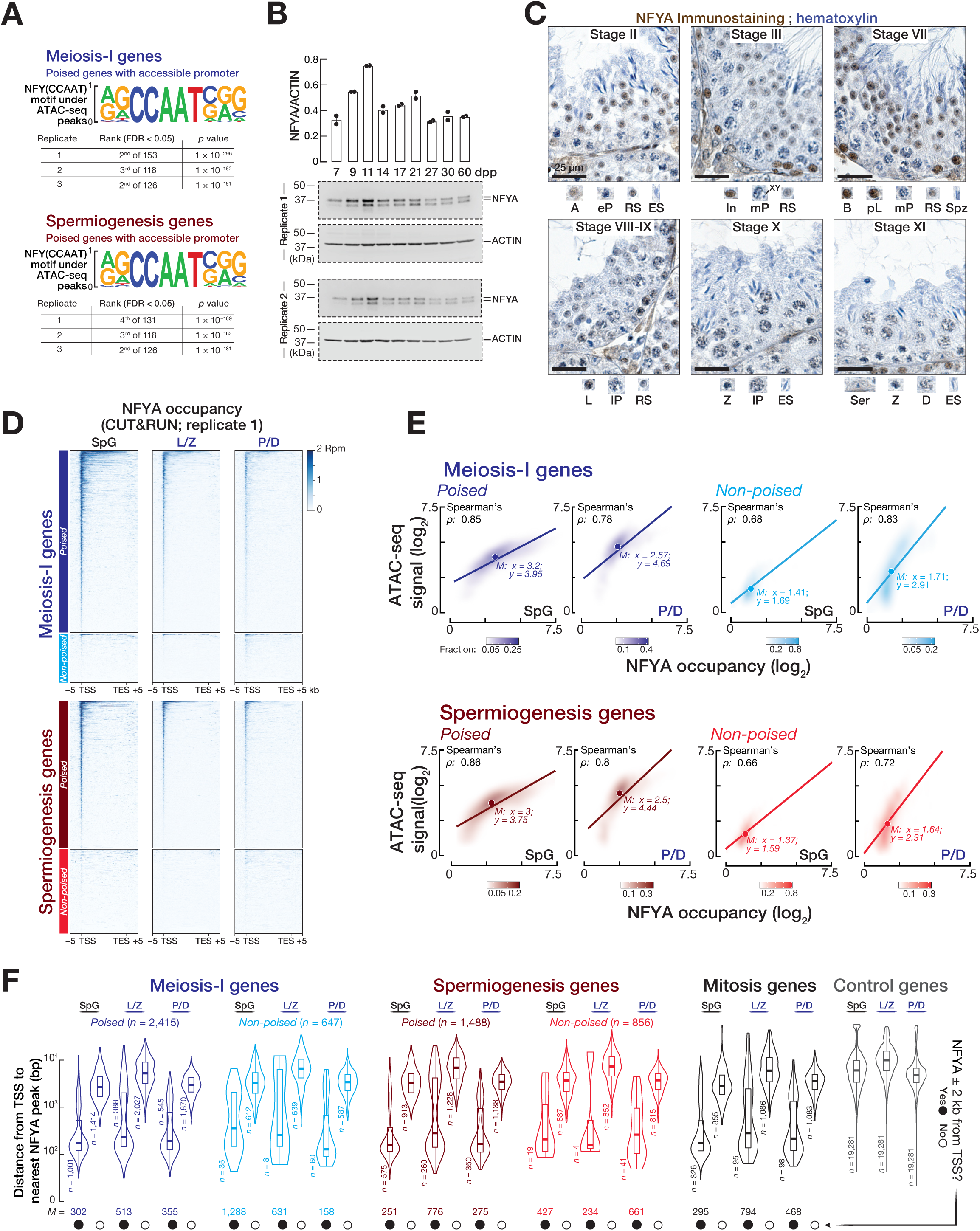
NFYA expressed highly in pre-meiotic cells binds the accessible promoters of poised genes in spermatogonia. **(A)** HOMER^59^ (Benjamini-Hochberg multiple test correction, FDR < 0.05) identified a sequence motif for NFYA under ATAC-seq peaks located around the promoters of poised genes (top: meiosis-I genes; bottom: spermiogenesis genes) in spermatogonia. Promoters of control genes (*n* = 19,281), whose transcript abundance remained constant across germ cells, served as background. **(B)** Abundance of NFYA protein across the testes of staged mice. ACTIN served as a loading control. Each lane contained 75 µg testis protein. Bars represent the mean protein abundance of *Nfya* from two independent replicates. Whiskers show standard deviation. Quantification of NFYA and ACTIN bands were performed using ImageJ2 v2.14. See Figure S5B for uncropped western blot images. **(C)** Immunohistochemical detection of NFYA in testis section from 3-month-old mouse. Seminiferous tubules at different epithelial stages are presented. A, type A spermatogonia; B, type B spermatogonia; In, intermediate spermatogonia; pL, preleptotene; L, leptotene; Z, zygotene; eP, early pachytene; mP, middle pachytene; lP, late pachytene; D, diplotene; RS, round spermatid; ES, elongating spermatid; Spz, spermatozoa; Ser, sertoli. **(D)** Heatmaps show reads per million (Rpm)-normalized NFYA CUT&RUN signals in the –5 kb to +5 kb window flanking transcription start sites (TSSs) and transcription end sites (TESs) of poised and non-poised genes for spermatogonia (SpG), leptotene/zygotene (L/Z), and pachytene/diplotene (P/D) cells. Heatmaps for second and third replicates are in Figure S4A. **(E)** Scatter plots show the spearman’s correlation (*P*) between ATAC-seq density and NFYA occupancy at the promoters of poised and non-poised meiosis-I and spermiogenesis genes. Each data point represents the average of three biological replicates. **(F)** Distance (average of three biological replicates) from the TSS to nearest NFYA peak for poised and non-poised meiosis-I and spermiogenesis genes, mitosis genes, and control genes in SpG, L/Z, and P/D cells. Genes were classified as NFYA-bound genes if they had significant NFYA peak ± 2 kb from their TSSs in at least two biological replicates of CUT&RUN experiment. Vertical lines represent median. Whiskers show maximum and minimum values. IQR is represented by boxplots. See also Figures S3, S5, and S6.

Consistent with the ubiquitous expression of NFYA^41^, mouse Encyclopedia of DNA Elements (ENCODE) revealed broad expression of *Nfya* mRNA across many mouse tissues (Figure S5A). To study the developmental expression of NFYA during spermatogenesis, we first measured protein abundance of *Nfya* across the testes of staged mouse (Figures 3B and S5B). By 9 to 11 days post-partum (dpp), the first cohort of spermatogenic cells enter prophase I and progress no further than zygotene cells. Hence, at 9 to 11 dpp, the majority of seminiferous tubules contain spermatogonia and early meiocytes of prophase I, leptotene and zygotene^42^. Intriguingly, NFYA abundance peaked at 11 dpp and was reduced in the testes of later stages (Figures 3B and S5B), suggesting that NFYA is expressed higher in spermatogonia or in early meiocytes compared to middle/late meiocytes and spermatids. To better assess the timing of NFYA expression during spermatogenesis, we immunostained the tubules with antibodies against NFYA (Figures 3C and S5C). To ensure robust conclusion from immunostaining, we used antibodies from two different companies (one is used for immunohistochemistry, the other is for immunofluorescence). Note that the specificity of antibodies used in this study was validated in *Nfya* mutant mice (Figures S7D and S7E). The cycle of mouse seminiferous epithelium is divided into 12 stages^43,44^, which provides an important map to understand the expression of a protein throughout germ cell development. In stage II tubules, type A spermatogonia moderately expressed NFYA, while tubules at stages III and VII exhibited a high abundance of nuclear NFYA protein in differentiating spermatogonia (i.e., intermediate and type B) and preleptotene spermatocytes that are the immediate descendants of spermatogonia appearing just before the entry into prophase I (Figure 3C). However, in the tubules at stages from VIII to XI, NFYA expression was gradually decreased from the leptotene stage to diplotene stages of prophase I (Figure 3C). Consistently, immunofluorescence staining revealed that those germ cells residing near basal membrane—intermediate and type B spermatogonia, as well as preleptotene spermatocytes—had the highest NFYA signal in their nuclei when compared to those localized more towards the lumen—leptotene, zygotene, pachytene, diplotene, and round spermatids (Figure S5C). Moreover, immunostaining of NFYA in the testes sections from staged mice corroborated these findings: we detected a strong NFYA signal localized in the nuclei of pre-meiotic cells, whilst the signal from meiocytes and spermatids was either low or below the limit of detection (Figure S5D). These results suggest that NFYA—expressed abundantly in pre-meiotic cells—may be a potential factor with a pioneer activity to promote accessible chromatin at poised gene promoters prior to meiotic entry.

### NFYA binds the promoters of poised genes in spermatogonia

To characterize the binding sites of NFYA genome-wide and to determine whether NFYA directly binds the promoters of poised genes in spermatogonia, we performed CUT&RUN from FACS-purified SpG, L/Z, and P/D cells using antibodies against NFYA and IgG. We sequenced three biological replicates for each cell population (moderate to strong agreement between replicates; SpG Spearman’s ρ 0.67–0.74, L/Z Spearman’s ρ 0.66–0.69, P/D Spearman’s ρ 0.57–0.63). We identified significant NFYA peaks, reflecting NFYA binding sites, genome-wide using MACS3 (FDR < 0.01; Table S3). IgG CUT&RUN served as control to map significant NFYA peaks. In SpG cells, 70% of all NFYA peaks were within the ±2 kb of annotated TSSs (Figure S6A). Consistently, in each SpG replicate, NFYA occupancy was higher at the TSSs compared with IgG control (Figure S6B). Moreover, only 2.2% of genome-wide NFYA peaks resided in enhancer regions marked by histone modifications, H3K4me1 (Figure S6A). Similar observations were obtained in L/Z and P/D cells (Figures S6A and S6B). Our analysis identified 4,020 protein-coding and 360 non-coding genes with an NFYA peak within ±2 kb of their TSSs in at least two replicates of SpG cells, whereas we found lesser number of genes bound by NFYA in L/Z and P/D cells consistent with the reduced expression of NFYA after meiotic entry (Figure S6C; Table S3). Together, we conclude that NFYA primarily functions at the promoter-proximal regions of genes.

We next sought to directly test that the promoters of poised genes expressed after meiotic entry are bound by NFYA in SpG cells. Remarkably, we detected strong NFYA occupancy around the TSSs of poised genes in SpG cells, whereas non-poised genes had no NFYA signal around their TSSs (Figures 3D, S4A, and S6D). Additionally, NFYA was expressed higher in pre-meiotic cells than in meiocytes (Figures 3B, 3C, S5B, and S5C). Consistent with this, NFYA occupancy at poised gene promoters in SpG cells (median = 9.1 rpm; median = 7.9 rpm; meiosis-I and spermiogenesis genes, respectively) was significantly higher than in L/Z (median = 4.7 rpm; median = 4.5 rpm; meiosis-I and spermiogenesis genes, respectively) and P/D cells (median = 5.9 rpm; median = 5.6 rpm; meiosis-I and spermiogenesis genes, respectively) (Figure S6E; Mann-Whitney-Wilcoxon U test). We next examined the correlation between NFYA occupancy and chromatin accessibility at the promoters of poised and non-poised genes in SpG and P/D cells. Strikingly, we found a strong correlation between NFYA occupancy and accessible chromatin at poised gene promoters (Figure 3E; Spearman’s ρ = 0.78–0.86). In fact, in SpG cells, the signal for both ATAC-seq and NFYA CUT&RUN at poised gene promoters was higher compared to non-poised gene promoters (Figure 3E). Note that although non-poised gene promoters displayed accessible chromatin in P/D cells, NFYA occupancy was not prominent suggesting that the establishment of permissive chromatin at non-poised gene promoters is likely through another factor whose expression perhaps appears at pachytene stage of prophase I (Figure 3E).

To determine the number of unambiguous NFYA-bound genes for each gene category, we computed the distance from the TSSs of poised and non-poised genes to the nearest NFYA peak in three biological replicates. Those genes with NFYA peak within the ±2 kb of their TSSs in at least two replicates were considered NFYA-bound genes. Notably, the promoters of ∼40% of poised genes were bound by NFYA in SpG cells (Figure 3F; 1,001 of 2,415 poised meiosis-I genes with median distance from TSS to the nearest NFYA peak = 302 bp; 575 of 1,488 poised spermiogenesis genes with median distance = 251 bp). On the contrary, only ≤5% of non-poised gene promoters were occupied by NFYA consistent with their closed chromatin state at SpG stage (Figure 3F; 35 of 647 non-poised meiosis-I genes; 19 of 856 non-poised spermiogenesis genes). Corroborating the reduced expression of NFYA after meiotic entry, the promoters of <25% of poised genes were occupied by NFYA in L/Z or P/D cells (Figure 3F). Among the 1,181 mitosis genes, NFYA bound to the promoters of 28% in SpG cells (Figure 3F; median distance = 295 bp). This suggests a regulatory function for NFYA in mitotic transcriptional program beyond its potential role in promoting accessible chromatin around the promoters of poised genes. Finally, examination of PRO-seq, ATAC-seq and NFYA CUT&RUN data on exemplified poised genes revealed an accumulation of paused Pol II at accessible promoters which is occupied by NFYA in SpG cells (Figure S6F). Our data suggests that prior to meiosis, NFYA may act as a pioneer factor to facilitate accessible chromatin at the promoters of poised genes that are expressed after meiotic entry.

### Male germ cell-specific deletion of *Nfya* severely impairs spermatogenesis

Prior to meiotic entry, NFYA—expressed highly in differentiating spermatogonia and preleptotene spermatocytes—binds the promoters of poised genes whose expression is in fact turned on at pachytene stage of prophase I. To understand the biological role of NFYA in meiotic entry and progression in a C57BL/6J background, we generated *Nfya^fl/fl^*; *Stra8-Cre* mice in which exons 5–6 of *Nfya* is deleted specifically in germ cells by the activity of Cre recombinase (Figures S7A–S7C; henceforth *Nfya-CKO*). *Stra8-Cre* mice expressed Cre recombinase from the promoter of *Stra8* gene (Figure S7A). Few days after birth, Stra8-Cre expression is active in undifferentiated and differentiating spermatogonia, and preleptotene spermatocytes^45^. Western blotting and immunohistochemistry analysis of staged mice testes confirmed the depletion of NFYA in germ cells, but not somatic cells (Figures S7D and S7E).

In *Nfya-CKO* male mice, we observed no abnormalities in organs other than testis. *Nfya-CKO* males had significantly smaller testis compared with control *Stra8-Cre^Cre/+^* males (henceforth *Cre/+*) (Figures S8A and S8B). Histological analysis of epididymis from 8-week-old mice additionally revealed a lack of sperm in *Nfya-CKO* males (Figure S8C). We thus conclude that *Nfya-CKO* males are infertile.

### NFYA-deficient germ cells arrest at meiotic entry

To gain understanding of the defective spermatogenesis in *Nfya-CKO* males, we histologically examined the testes at 9, 14, and 30 dpp (Figure 4A). In *Cre/+* mice, by 9 dpp, the first cohort of spermatogenic cells reached leptotene cells; at 14 dpp, early pachytene spermatocytes appeared; while in 30 dpp, spermatogenesis reached to spermatozoa (Figure 4A). Notably however, in *Nfya-CKO* males, spermatogenesis progressed no further than spermatocytes resembling preleptotene spermatocytes (Figure 4A). We next immunostained the tubules for DDX4, a marker for both male and female germ cells^46^. Here, by 9 dpp, spermatogenesis reaches no later than leptotene cells—accounting for the smallest fraction of all germ cells^42^. Hence, we observed a comparable number of DDX4-positive germ cells between *Nfya-CKO* and *Cre/+* mice at 9 dpp (Figure S8D). However, at 14 and 30 dpp, DDX4-positive germ cells were markedly reduced in *Nfya-CKO* males when compared with *Cre/+* mice (Figure S8D). In the testis sections from *Nfya-CKO* males, we observed germ cells displaying morphological characteristics of dying cells (Figure 4A). Subsequently, we sought to examine the fate of defective germ cells in *Nfya-CKO* by performing the terminal dUTP nick-end labeling (TUNEL) staining. We found that the seminiferous tubules of *Nfya-CKO* testis had more TUNEL-positive germ cells compared to *Cre/+* testis, suggesting that defective germ cells in *Nfya-CKO* testis may be eliminated in part by apoptosis (Figure S8E).

**Figure 4.**
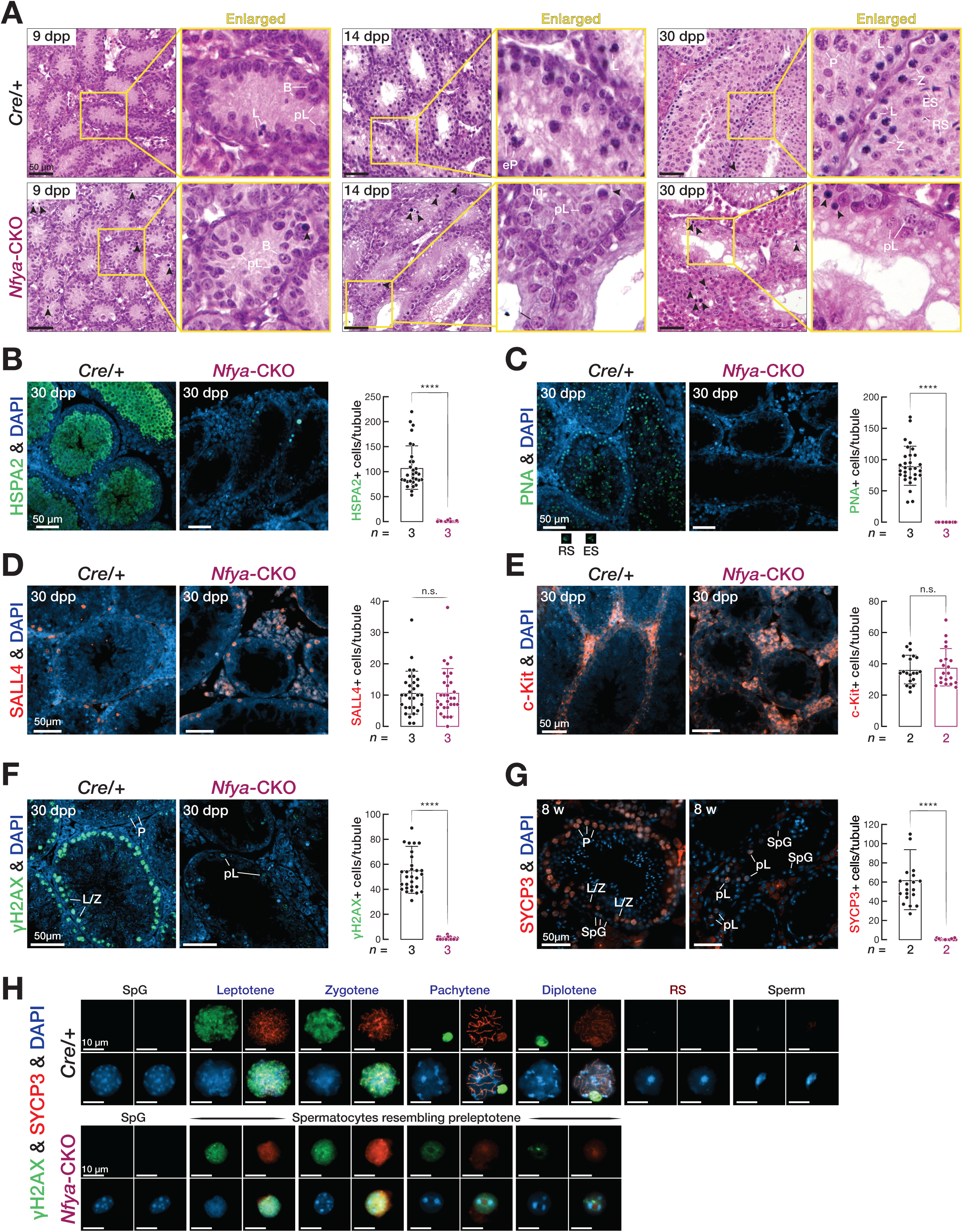
Germline-conditional deletion of *Nfya* blocks meiotic entry. **(A)** Hematoxylin and eosin staining of the sections from *Cre/+* and *Nfya-CKO* mice at 9 dpp, 14 dpp, and 30 dpp. Right: Enlarged images. Black arrow heads depict dying cells. B, type B spermatogonia; In, intermediate spermatogonia; pL, preleptotene; L leptotene; Z, zygotene; eP, early pachytene; P, pachytene; RS, round spermatid; ES, elongating spermatid. Scale bars: 50 μm. **(B-G)** Immunofluorescence staining of the sections from 30 dpp and 8-week-old *Cre/+* and *Nfya-CKO* mice for **(B)** HSPA2, **(C)** Peanut agglutinin (PNA), **(D)** SALL4, **(E)**, c-Kit **(F)** γH2AX, and **(J)** SYCP3. All images are representative of two or three biological replicates. Bars represent quantification for positive cells per tubule from two or three independent biological replicates for each staining. Whiskers show standard deviation. Significance is measured using two-sided unpaired student t-test. Scale bars: 50 μm. **(H)** Immunofluorescence staining for γH2AX and SYCP3 from the chromosome spreads of *Cre/+* and *Nfya-CKO* germ cells. Scale bars: 10 μm. See also Figures S7 and S8.

Spermatogenic defect appears to occur at meiotic entry rather than during meiotic progression, as judged from the morphology of germ cells which resemble preleptotene spermatocytes (Figure 4A). To substantiate this observation, we immunostained the tubules with antibody against HSPA2, testis-specific member of HSP70 family that is expressed from pachytene spermatocytes and onward^47^. We found that HSPA2-positive pachytene spermatocytes and germ cells were absent in *Nfya-CKO* testis (Figure 4B). Moreover, immunostaining for peanut agglutinin (PNA), a plant lectin that binds the contents of acrosomes in developing spermatids and spermatozoa^48^, further confirmed the absence of round and elongating spermatids in *Nfya-CKO* testis (Figure 4C). Together, our data demonstrates that NFYA deficient spermatocytes—that resemble preleptotene cells—fail to progress through prophase I.

In *Nfya-CKO* testis, spermatogenesis appears to arrest at the preleptotene stage. To test the idea that the observed defect in spermatogenesis occurs during meiotic entry rather than during the maintenance of undifferentiated spermatogonia or during the differentiation of spermatogonia into spermatocytes, we immunostained the tubules for SALL4—a marker for undifferentiated spermatogonia^26^—and c-Kit, a marker for differentiating spermatogonia^49^. Here, we found a comparable number of SALL4-positive undifferentiated spermatogonia and c-Kit-positive differentiating spermatogonia between *Nfya-CKO* and *Cre/+* mice (Figures 4D and 4E). These findings suggests that the maintenance and differentiation of spermatogonia occur normally within *Nfya-CKO* testis.

Formation of DNA double-strand break (DSB) and waves of expression of components of axial elements (AE) are hallmarks of meiotic initiation^5^. We thus performed immunostaining for γH2AX, a marker for DSB, and SYCP3, a component of AE, to directly test that NFYA-deficient germ cells reach preleptotene stage but fail to progress through prophase I. γH2AX staining demonstrated the presence of DSBs in a greater number of meiocytes from *Cre/+* testis when compared to those from *Nfya-CKO* testis. Nonetheless, we detected a γH2AX signal in the nuclei of spermatocytes resembling preleptotene cells (Figures 4F and S8F). Moreover, the nuclei of spermatocytes resembling preleptotene cells additionally showed a SYCP3 signal (Figure 4G). To further substantiate these observations, we performed double staining for γH2AX and SYCP3 on chromosome spread from the cells of *Nfya-CKO* and *Cre/+* testis. While germ cells successfully progressed through meiosis— as evidenced by the presence of leptotene, zygotene, pachytene and diplotene spermatocytes—and developed into spermatozoa in *Cre/+* testis, spermatocytes later than the preleptotene stage did not appear in *Nfya-CKO* testis (Figures 4H). Notably, spermatocytes resembling preleptotene cells showed a fuzzy and aggregated SYCP3 signal, suggesting incomplete AE formation (Figures 4H). Together, these results demonstrate that NFYA-deficient germ cells developed into preleptotene spermatocytes, but stalled there to progress through meiosis.

### Simultaneous scRNA-seq and scATAC-seq reveals lack of meiocytes in *Nfya-CKO*

To directly test whether the arrest at meiotic entry in *Nfya-CKO* mice is due to the disruption of chromatin accessibility at meiotic promoters in spermatogonia, we performed simultaneous single-cell RNA-seq (scRNA-seq) and single-cell assay for transposase accessible chromatin sequencing (scATAC-seq) on the nuclei from the testes of 8-week-old *Cre/+* and *Nfya-CKO* mice, as well as 11, 17, and 25 dpp wild-type mice using BD Rhapsody Single-Cell Analysis System. For scRNA-seq, a total of 45,250 individual cells were captured. We detected an average of 1,977 genes per cell and an average of 5,850 copies of transcripts (unique molecular indices, UMIs) per cell (Table S4A). We performed uniform manifold approximation and projection (UMAP) using Seurat package^50^ for dimension reduction analysis on the combined datasets. Independent of their origin of dataset, cells formed 32 clusters indicating minimal batch effect (Figure S9A). Using previously described cell-type specific markers (Figure S9B; Table S4B), we further resolved 32 clusters into 13 cell populations: 9 major germ cell populations—undifferentiated spermatogonia (undiff. SpG), differentiating spermatogonia (diff. SpG), preleptotene spermatocytes (pL), early leptotene, leptotene, zygotene spermatocytes (eL/L/Z), leptotene, zygotene, early pachytene spermatocytes (L/Z/eP), middle and late pachytene spermatocytes (mP/lP), diplotene spermatocytes (D), round spermatids (RS), and elongating spermatids (ES)—; two cell populations from *Nfya-CKO* mice—spermatogonia and somatic cells—; Sertoli cells; and unknown cells (Figures 5A–5C).

**Figure 5.**
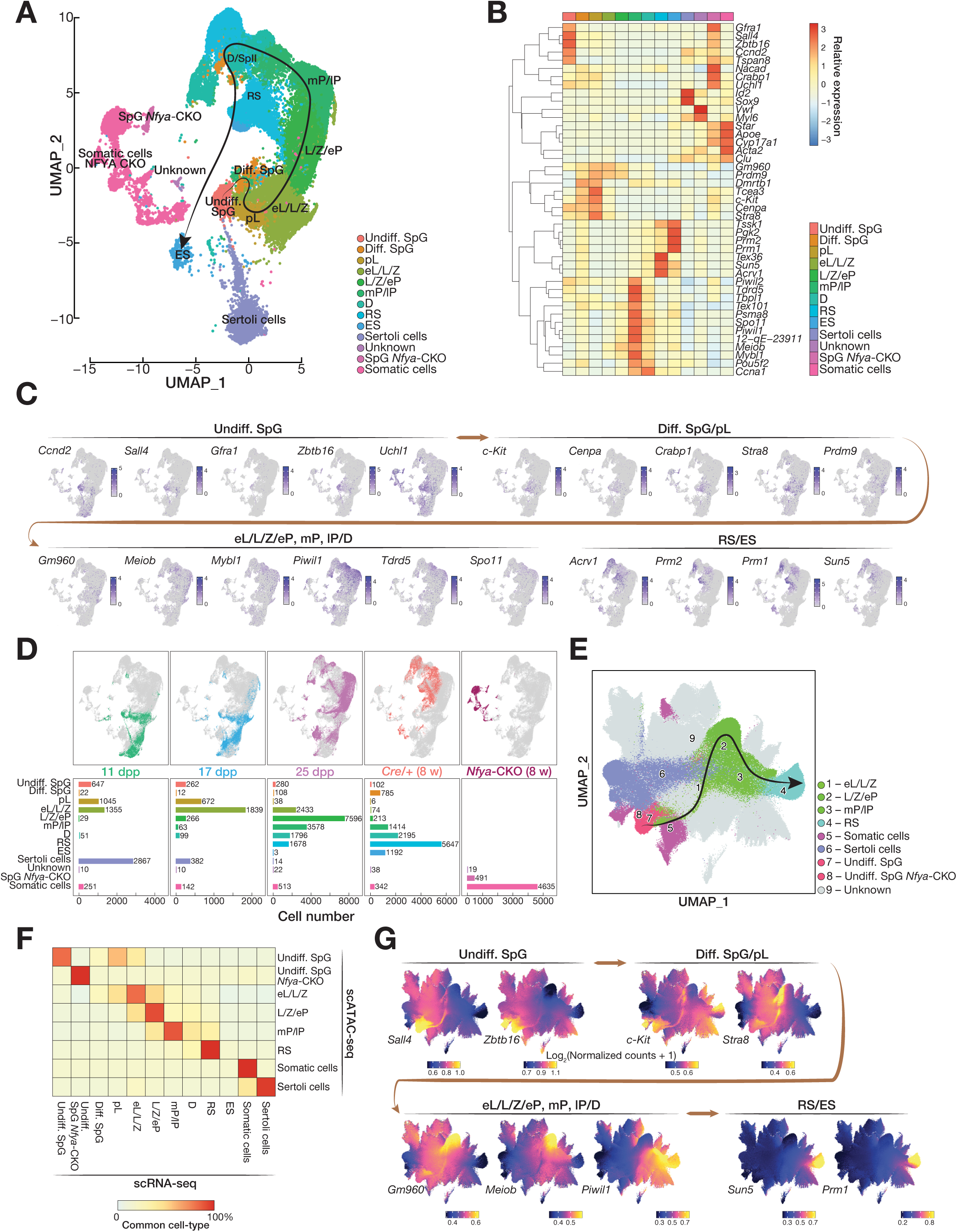
Identifying cellular diversity of testes from 8-week-old *Cre/+* and *Nfya-CKO* mice along with wild-type staged mice using simultaneous single-cell (sc) RNA-seq and ATAC-seq. **(A)** Dimension reduction analysis by uniform manifold approximation and projection (UMAP) on 45,250 individual cells captured from all datasets illustrates 13 major cell populations in different colors. Major cell populations are defined from 32 clusters using the spatiotemporal expression of previously described cell-type specific marker genes. See Figure S9A for the full set of 32 clusters. Undiff. SpG, undifferentiated spermatogonia; Diff. SpG, differentiating spermatogonia; pL, preleptotene; eL/L/Z, early leptotene, leptotene, and zygotene; L/Z/eP, leptotene, zygotene, early pachytene; mP/lP, middle and late pachytene; D, diplotene; RS, round spermatid; ES, elongating spermatid; Sertoli cells; Unknown cells; SpG *Nfya-CKO*, spermatogonia from *Nfya-CKO*; other somatic cells. **(B)** Heatmap illustrates the relative expression levels of all marker genes included in the study across 13 annotated cell populations. **(C)** Per-cell gene expression patterns for exemplified known cell-type specific marker genes are visualized in UMAP space. Transitions between major cellular states are highlighted in brown color. **(D)** UMAP spaces (top) illustrate the identity of cellular diversity in each mouse compared to the background of remaining mice (grey color). Bars (bottom) represent the total number of individual cells belonging to each major cell population defined in panel A in each mouse. **(E)** UMAP space visualizes nine cell populations as defined by the chromatin accessibility state of 343,866 individual cells obtained from scATAC-seq. Major cell populations are defined from the 25 clusters by the guidance from scRNA-seq based cell-type annotation as well as the chromatin state of marker genes. See Figure S9C for the full set of 25 clusters. **(F)** Heatmap shows the percentage of cells that are classified in the same cell-type in both scRNA-seq and scATAC-seq. **(G)** Dynamic change in permissive chromatin state around the promoters of exemplified known cell-type specific marker genes in individual cell is illustrated in UMAP space. Transitions between major cellular states are highlighted in brown color. See also Figure S9.

Differences for cell-type composition across the testes from staged and *Cre/+* and *Nfya-CKO* mice recapitulated both the timeline of germ cell development throughout stages and our observed phenotype for *Nfya-CKO* mice (Figure 5D). Developing germ cell populations agreed well with the expected developmental stage^42^. At 11 dpp, the most developed germ cell population was eL/L/Z, whereas 17dpp reached L/Z/eP. By 25 dpp, we detected all kinds of spermatocytes, as well as haploid spermatids. Testis of 8-week-old *Cre/+* contained entire germ cell populations, while cell populations from *Nfya-CKO* mice occupied distinct locations in UMAP space, reflecting the absence of spermatocyte populations. In fact, the major cell populations were somatic cells and spermatogonia, consistent with the spermatogenic arrest at meiotic entry observed in *Nfya-CKO* mice (Figure 5D).

We next analyzed the scATAC-seq dataset of our simultaneous sc-multiomics experiment. After filtering low-quality cells and doublets, we captured 343,866 cells with high quality accessible chromatin for downstream analysis (average TSS enrichment score per cell = 11.3; Table S4C). Through clustering based on a genome-wide tile matrix of 500 bp using ArchR^51^ on the combined cells from the testes of both 8-week-old *Cre/+* and *Nfya-CKO* mice, as well as 11, 17, and 25 dpp wild-type mice, our analysis identified 25 clusters (Figure S9C). To annotate the cell types from 25 clusters, we used our scRNA-seq based cell-type annotation for cells with both accessible chromatin and captured transcriptome (Figure S9D). Here, our analysis resolved 25 clusters into 9 cell populations (eL/L/Z, L/Z/eP, mP/lP, RS, somatic, Sertoli, undiff. SpG, undiff. SpG from *Nfya-CKO*, and unknown) which agreed well with the cell populations obtained from scRNA-seq (Figures 5E and 5F). Determination of chromatin accessibility around the TSSs of key cell-type specific marker genes further corroborated the annotation of major cell populations (Figure 5G). Finally, cell-type composition differences in each dataset recapitulated the developmental trajectory of germ cell and the spermatogenic arrest in *Nfya-CKO* mice (Figure S9E). the testes of wild-type staged and 8-week-old *Cre/+* mice, we observed step-wise development of male germ cells. Whereas, in 8-week-old *Nfya-CKO* testes, the major cell populations we captured were somatic cells and spermatogonia (Figure S9E). In summary, our systematic analysis of simultaneous scRNA-seq and scATAC-seq demonstrated continuous development of germ cells in wild-type and *Cre/+* testes, and provided further supporting evidence that spermatogenesis arrests at meiotic entry in *Nfya-CKO* mice.

### *Nfya* deletion impairs chromatin accessibility at the promoters of poised genes in spermatogonia

We defined three gene categories based on the change in absolute transcript abundance of genes across four major FACS-purified bulk germ cell populations (Figure S1A). Supporting our classification, unsupervised hierarchical clustering based on the expression of mitosis, meiosis-I, and spermiogenesis genes from nine cell populations captured by scRNA-seq revealed their temporal expression across spermatogenesis (Figure 6A). The expression of mitosis genes was highest in undifferentiated spermatogonia and sharply reduced in early meiocytes (i.e., leptotene/zygotene/early pachytene). In contrast, transcripts of meiosis-I genes first accumulated in early meiocytes (i.e., early leptotene/leptotene/zygotene); peaked in middle and late pachytene; and substantially declined in round spermatids. The expression of spermiogenesis genes coincided with meiosis-I genes yet persisted in spermatids (Figures 6A and 6B).

**Figure 6.**
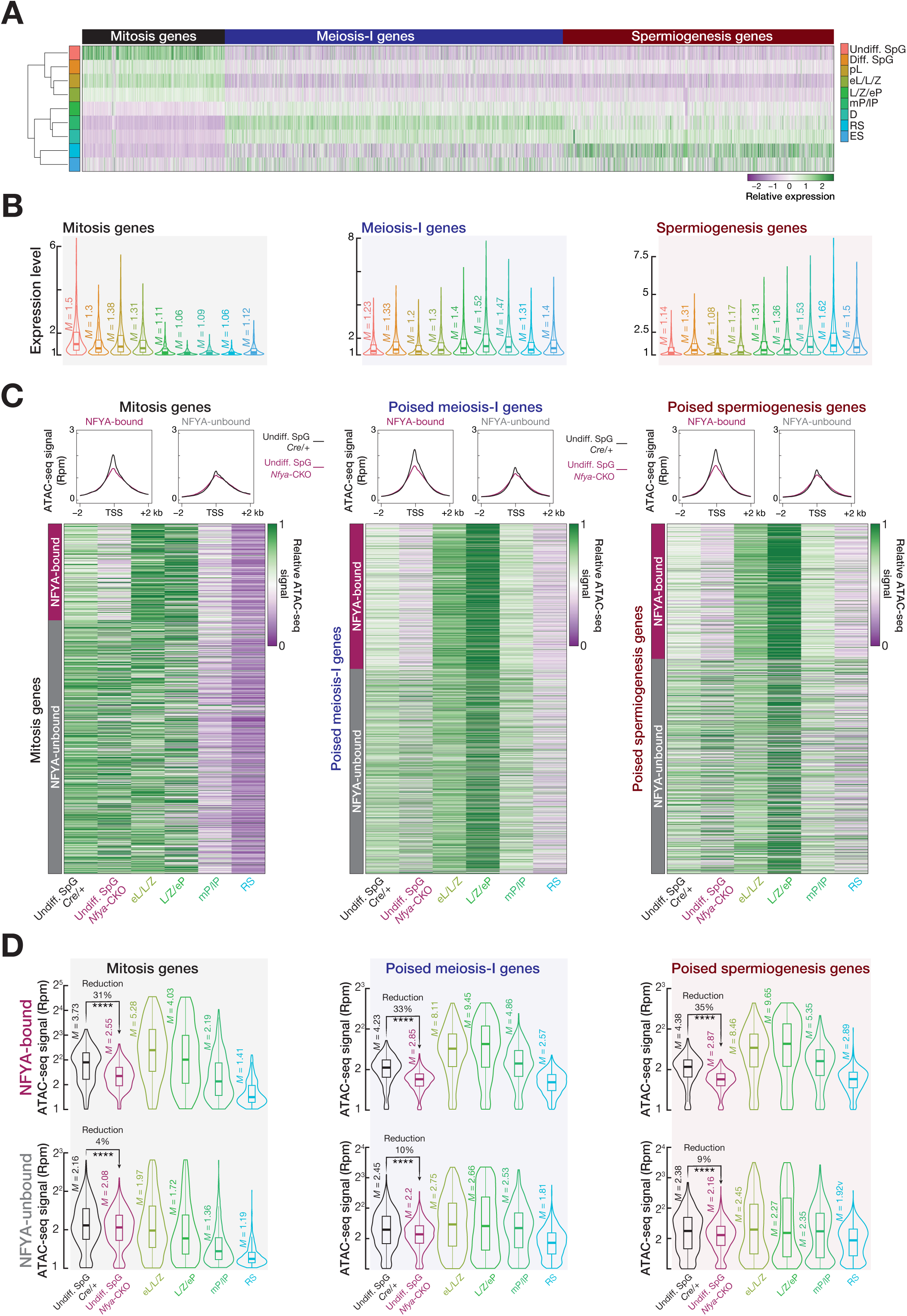
*Nfya* deletion impairs chromatin accessibility at meiotic gene promoters in spermatogonia. **(A)** Unsupervised hierarchical clustering of nine germ cell populations captured by sc-RNA using the transcript abundance of mitosis, meiosis-I, and spermiogenesis genes. Undiff. SpG, undifferentiated spermatogonia; Diff. SpG, differentiating spermatogonia; pL, preleptotene; eL/L/Z, early leptotene, leptotene, and zygotene; L/Z/eP, leptotene, zygotene, early pachytene; mP/lP, middle and late pachytene; D, diplotene; RS, round spermatid; ES, elongating spermatid. **(B)** RNA expression levels of mitosis, meiosis-I and spermiogenesis genes across nine germ cell populations. Vertical lines represent median. Whiskers show maximum and minimum values. IQR is represented by boxplots. **(C)** Metaplot (top) shows ATAC-seq signal in the –2 kb to +2 kb window flanking transcription start sites (TSSs) of NFYA-bound and -unbound mitosis, poised meiosis-I and poised spermiogenesis genes. Heatmap (bottom) represents relative ATAC-seq signal at the promoters of mitosis, poised meiosis-I and poised spermiogenesis genes that are either NFYA-bound or -unbound. **(D)** Reads per million (Rpm)-normalized ATAC-seq signals at the promoters of mitosis, poised meiosis-I, and poised spermiogenesis genes which are either NFYA-bound or - unbound. Vertical lines represent median. Whiskers show maximum and minimum values. IQR is represented by boxplots. See also Figure S9.

Prior to the initiation of meiosis, in pre-meiotic cells, NFYA occupies the promoters of genes that remain poised for activation later during meiosis. To directly test the idea that NFYA regulates chromatin accessibility in pre-meiotic cells, we examined whether *Nfya* deletion disturbed chromatin accessibility at meiotic promoters in spermatogonia. We analyzed changes in chromatin accessibility around the promoters of NFYA-bound and -unbound gene categories (Figure 3F) in undifferentiated spermatogonia from *Cre/+* and *Nfya-CKO* mice. Remarkably, we found that relative scATAC-seq signal at the promoters of NFYA-bound poised meiosis-I and spermiogenesis genes were reduced in NFYA-deficient undifferentiated spermatogonia compared to that from *Cre/+* mice (Figure 6C; Table S4D). In contrast, ATAC-seq signal at the promoters of NFYA-unbound poised genes remained almost unchanged between undifferentiated spermatogonia from *Cre/+* and *Nfya-CKO* mice (Figure 6C; Table 4D). In fact, chromatin accessibility at the promoters of NFYA-bound poised genes were reduced ∼35% in undifferentiated spermatogonia from *Nfya-CKO*, whereas the reduction for NFYA-unbound poised genes was ∼10% in NFYA-deficient undifferentiated spermatogonia (Figure 6D; Two-sided wilcoxon matched-pairs signed ranked sum test; \*\*\*\**p* < 0.0001). As we did not observe phenotypic perturbation in spermatogonia from *Nfya-CKO* mice (Figure 4), the observed change in promoter accessibility of NFYA-bound mitosis genes (Figure 6C and D) can perhaps be tolerable for cells to progress through spermatogonia differentiation. Alternatively, those genes with reduced accessibility are not necessary for spermatogonia differentiation. Indeed, promoter accessibility of key factors required for spermatogonia maintenance and differentiation—e.g., *Sall4*, *Zbtb16*, *Nanos3*, *Thy1*, and *Kitl*—was unchanged in *Nfya-CKO* (Table S4D). Excluding *Sall4*, the promoters of these genes are not bound by NFYA at their promoters (Table S3). When taken together, we thus conclude that prior to meiotic entry, NFYA regulates chromatin accessibility at the promoters of genes that remain poised for activation later during meiosis.

### NFYA establishes a coherent feedforward loop to regulate STRA8/MEIOSIN and their targets

TFs STRA8 and MEIOSIN induce the transition from mitosis to meiosis^10,11,52^. Similar to the defective phenotype reported in *Stra8*^10^ and *Meiosin*^11^ mutant mice, spermatogenesis arrests at meiotic entry in *Nfya-CKO* mice. In fact, we identified NFYA peaks around the promoters of *Stra8* and *Meiosin* genes in all three replicates of NFYA CUT&RUN from spermatogonia (Figure 7A; Table S3). Chromatin accessibility around the promoters of *Stra8* and *Meiosin* was reduced in undifferentiated spermatogonia from *Nfya-CKO* mice compared to that from *Cre/+* mice (Figure 7A; ∼10% and ∼30% reduction for *Stra8* and *Meiosin*, respectively).

**Figure 7.**
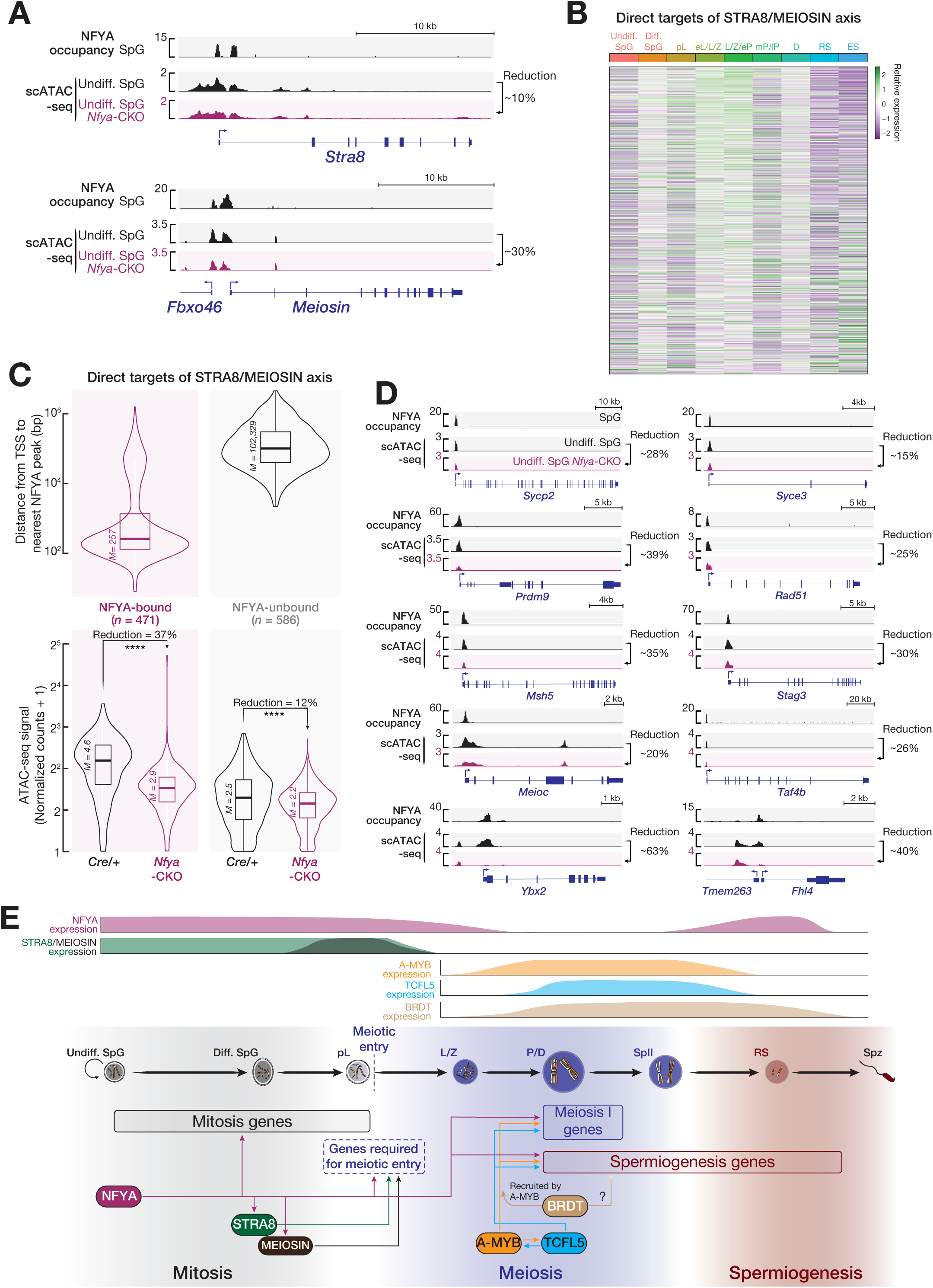
NFYA regulates chromatin accessibility at the promoters of *Stra8* and *Meiosin* genes as well as STRA8/MEIOSIN target genes via a coherent feedforward loop. **(A)** Integrative Genomics Viewer (IGV) views for NFYA CUT&RUN and ATAC-seq signals at gene boundaries of *Stra8* (top) and *Meiosin* (bottom). **(B)** Heatmap shows relative expression of genes targeted by STRA8/MEIOSIN axis in nine defined populations from scRNA-seq. Undiff. SpG, undifferentiated spermatogonia; Diff. SpG, differentiating spermatogonia; pL, preleptotene; eL/L/Z, early leptotene, leptotene, and zygotene; L/Z/eP, leptotene, zygotene, early pachytene; mP/lP, middle and late pachytene; D, diplotene; RS, round spermatid; ES, elongating spermatid. **(C)** Distance from transcription start site (TSS) to nearest NFYA peak (top) of genes targeted by STRA8/MEIOSIN axis. ATAC-seq signal (bottom) around the TSSs of genes targeted by STRA8/MEIOSIN axis in undifferentiated spermatogonia from *Cre/+* and *Nfya-CKO*. Vertical lines represent median. Whiskers show maximum and minimum values. IQR is represented by boxplots. **(D)** Integrative Genomics Viewer (IGV) views for NFYA CUT&RUN and ATAC-seq signals at gene boundaries of exemplified genes targeted by STRA8/MEIOSISN axis. **(E)** The model incorporates findings from refs.^10,11,13,14,17,19,20,31,52^ and this study. Germ cell development is aligned with the temporal expression of key transcription factors. The model highlights the sequential roles of key regulatory proteins, i.e, NFYA, STRA8, MEIOSIN, A-MYB, TCFL5, BRDT, during spermatogenesis. See also Figure S9.

We next sought to examine the change in chromatin accessibility at the promoters of 1,057 genes whose transcription is directly regulated by both STRA8 and MEISON or STRA8 alone, or MEIOSIN alone^11,52^. In fact, RNA abundance of many STRA8/MEIOSIN target genes increased in early meiocytes of prophase I (Figure 7B). Intriguingly, of the 1,057 genes, the promoters of 471 were bound by NFYA in spermatogonia (Figure 7C; top panel; median distance from TSS to the nearest NFYA peak = 257 bp). We found that chromatin accessibility around the promoters of NFYA-bound STRA8/MEIOSIN target genes was reduced ∼37% in undifferentiated spermatogonia from *Nfya-CKO* mice when compared to that from *Cre/+* mice (Figure 7C; bottom panel; Two-sided wilcoxon matched-pairs signed ranked sum test; \*\*\*\**p* < 0.0001). Whereas chromatin accessibility around the promoters of STRA8/MEIOSIN targets, which were not bound by NFYA, was nearly unchanged (Figure 7C; bottom panel). Among those 471 NFYA-bound STRA8/MEIOSIN target genes whose promoter accessibility was reduced in NFYA-deficient undifferentiated spermatogonia, many function in meiotic recombination (e.g., *Sycp2*, *Syce3*, *Prdm9*, *Rad51*, *Msh5*, *Stag3*^1,53^), cell cycle (e.g., *Meioc*^54^), and meiotic transcriptional program (e.g., *Taf4b*^55^) (Figure 7D). Taken together, our data supports a model in which NFYA facilitates accessible chromatin at the promoters of both *Stra8*/*Meiosin* and their target genes via a coherent feedforward loop during the transition from mitosis to meiosis (Figure 7E).

## DISCUSSION

Entry into the first and longest stage of meiosis, prophase I, accompanies extensive cellular and chromosomal events^1,2,5^. Such dramatic changes upon meiotic entry entails synchronous transcriptional activation of genes during pachytene stage of prophase I^13,14,17,20^. Importantly, in pre-meiotic spermatogonia, the promoters of many of these genes accumulate paused Pol II that is later released into elongation in pachytene stage of prophase I by the activity of TFs A-MYB and BRDT^19,20^. Prior to our findings reported here, how chromatin accessibility is established to allow Pol II to bind the promoters of meiotic genes in pre-meiotic cells was unknown. Here, we provide functional and molecular evidence that NFYA acts as a pioneer factor in pre-meiotic cells to facilitate chromatin accessibility around the promoters of genes expressed during prophase I. Intriguingly, our data also reveal that during meiotic entry NFYA promotes the promoter accessibility of two key meiotic initiators STRA8 and MEIOSIN^10,11,52^, and their target genes, thereby establishing a coherent feedforward loop. Together with previous findings^10,11,13,14,17,19,20,31,52^, our data suggests a model for a TFs network of spermatogenesis, in which NFYA plays a pivotal role by regulating the chromatin accessibility required during both meiotic entry and progression of prophase I (Figure 7E).

Given that NFYA binds the promoters of ∼40% of genes that are not yet expressed in spermatogonia (Figure 3) and that the transcription of many of these genes are later driven by a master TF, A-MYB, in pachytene spermatocytes (Figure S2), our findings raise an attractive possibility that NFYA functions as a pioneer factor in pre-meiotic cells. Supporting this possibility, chromatin accessibility at the promoters of NFYA-bound genes expressed during meiosis were markedly reduced in NFYA-deficient spermatogonia (Figure 6). In mouse ESCs, NFYA promotes chromatin accessibility for the binding of master TFs^56^. In fact, a model proposes that NFYA binding dictates ∼80% bend in DNA, ultimately resulting in nucleosome repositioning^56^. As another *in vitro* study demonstrated that NFYA displaces nucleosomes^57^, these findings suggest that NFYA retains intrinsic chromatin remodeling activity. However, we cannot rule out the possibility that NFYA recruits a chromatin remodeler to promote permissive chromatin. It will thus be an important subject of investigation in our future studies.

STRA8 and its partner MEIOSIN initiate meiotic entry by regulating both genes required for chromosomal events of prophase I as well as genes required for meiotic G1-S cell cycle regulation^11,52^. Spermatogenic arrest occurred at meiotic initiation rather than meiotic progression in *Stra8* and *Meiosin* mutant mice^10,11^. However, ectopic expression of STRA8 or MEIOSIN did not suffice for the induction of meiosis *in vitro*^11,58^, suggesting that meiotic entry requires additional molecular events acting upstream of STRA8/MEIOSIN. We would thus argue that the permissive chromatin state at the promoters of genes required for meiotic entry is established in pre-meiotic spermatogonia that precedes the activity of STRA8/MEIOSIN. Supporting this idea, our data revealed that the occupancy of NFYA at *Stra8*/*Meiosin* and many of their target gene promoters whose chromatin accessibility was reduced in NFYA-deficient spermatogonia (Figure 7). Indeed, we observed a lesser reduction in chromatin accessibility at the promoter of *Stra8* than that of *Meiosin* suggesting the existence of another factor compensating for NFYA activity. Because STRA8 and NFYA were expressed earlier than MEIOSIN (Table S4), it is possible that the expression of this compensatory factor precedes NFYA expression.

In summary, we have elucidated the role of NFYA in spermatogenesis. Specifically, we have shown that NFYA functions as a pioneer factor before meiotic entry to facilitate permissive chromatin around the promoters of genes required for meiotic entry, meiotic progression and spermiogenesis, thereby priming them for timely expression during spermatogenesis. Therefore, our discovery will benefit both our understanding of the stepwise development of male germ cells and the efforts for generating *in vitro*-derived germ cells for therapeutic purposes.

### Limitations of the study

Our data reveal that NFYA promotes accessible chromatin at poised gene promoters in spermatogonia and that chromatin accessibility is impaired in NFYA-deficient spermatogonia. We believe that permissive chromatin state in turn enables the accumulation of paused Pol II at poised promoters in spermatogonia. However, the requirement of a dazzling number of cells for PRO-seq limits us to directly measure the change in engaged paused Pol II at poised gene promoters in NFYA-deficient spermatogonia.

Although the initiation and the progression of meiosis is sexually dimorphic, the molecular mechanisms, such as the role of STRA8 and MEISOIN, are conserved between males and females. As NFYA is broadly expressed in many tissues including ovary (Figure S5A), it would be interesting to examine whether the pioneering role of NFYA is conserved in oocytes.

## Supporting information

Legends for supplemental figures

## RESOURCE AVAILABILITY

### Lead contact

Further information and requests for resources and reagents should be directed to and will be fulfilled by the Lead Contact, Deniz M. Özata.

### Data and code availability

Sequencing data are available from the National Center for Biotechnology Information Sequence Read Archive using Gene Expression Omnibus (GEO) accession number GSE295991 (see the Key resources table). All publicly available data were downloaded from GEO and are summarized in the Key resources table. All Scripts used to analyze sequencing data can be found here: https://github.com/Ozatalab. Original western blotting images are deposited at Mendeley Data, V1, doi: 10.17632/bycn6xz4tt.1, and are publicly available as of the date of publication. Microscopy data reported in this paper will be shared by the lead contact upon request. Any additional information required to reanalyze the data reported in this paper is available from the lead contact upon request.

## ACKNOWLEDGEMENTS

We thank the personnel of Stockholm University animal facility, with particular gratitude to S. Oerther and R. Askar for their excellence in mouse colony management; C. Molenaar for his expert help with Zeiss Axio Observer 7 inverted widefield microscope; National Genomics Infrastructure—funded by Science for Life Laboratory, the Knut and Alice Wallenberg Foundation and the Swedish Research Council—, and SNIC/Uppsala Multidisciplinary Center for assistance with massively parallel sequencing and access to the UPPMAX computational infrastructure. This work was supported by the Swedish Research Council grant 2020-03818 (D.M.Ö), the Swedish Research Council grant 2024-04321 (D.M.Ö), and Carltryggersstiftelse CTS 21:1158 (D.M.Ö).

## AUTHOR CONTRIBUTIONS

D.M.Ö. conceived and designed the experiments. M.S., M.A., A.E., and M.M.A. performed the experiments. M.A., and Y.T.X. analyzed the sequencing data. M.S., A.E., M.M.A., and D.M.Ö. characterized the *Nfya-CKO* mice phenotype. M.S., M.A., A.E., AK.I.Ö.F., Y.T.X., and D.M.Ö. wrote the manuscript. All authors discussed and approved the manuscript.

## DECLATION OF INTERESTS

The authors declare no competing interests.

## STAR⍰METHODS

### KEY RESOURCES TABLE

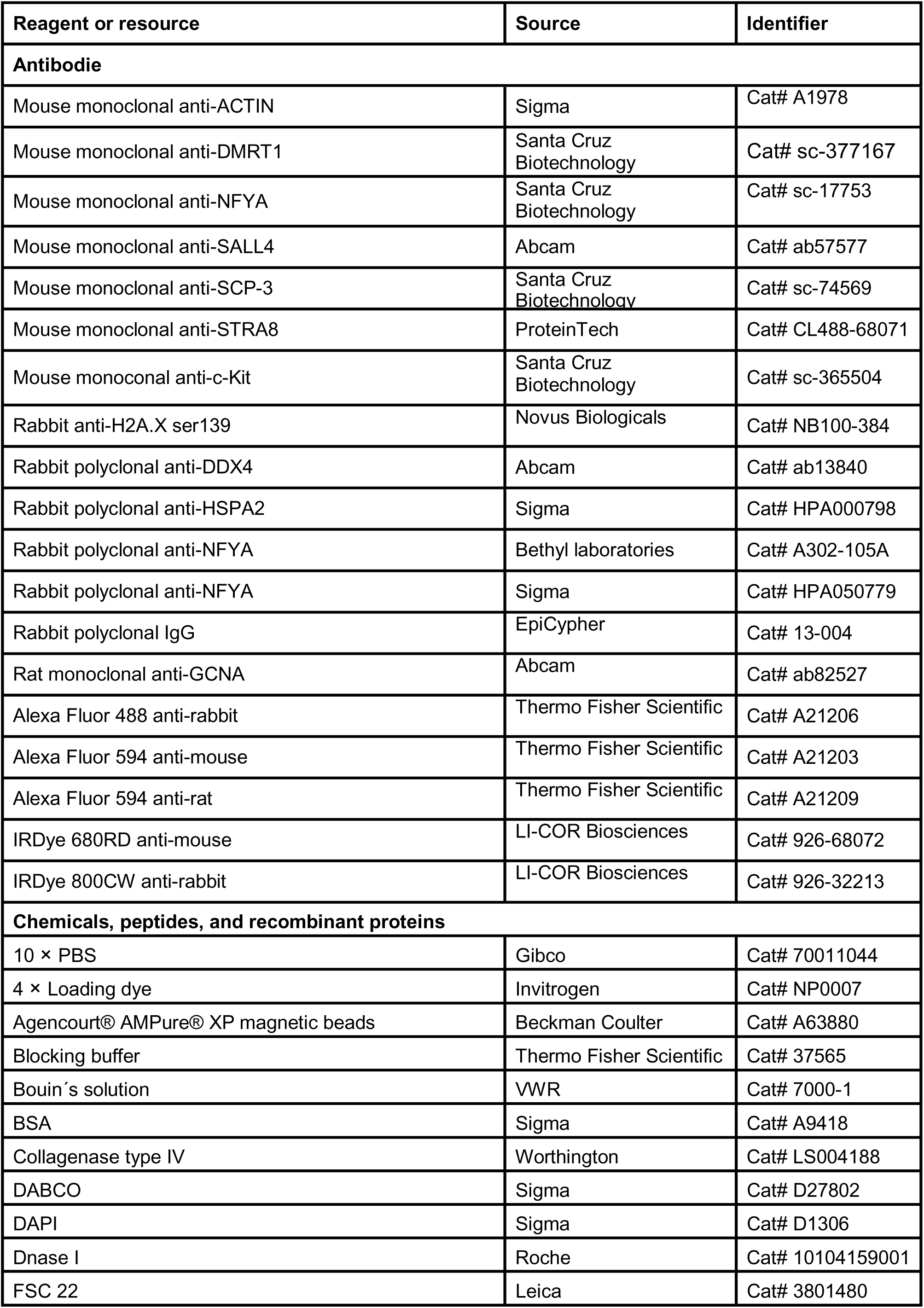
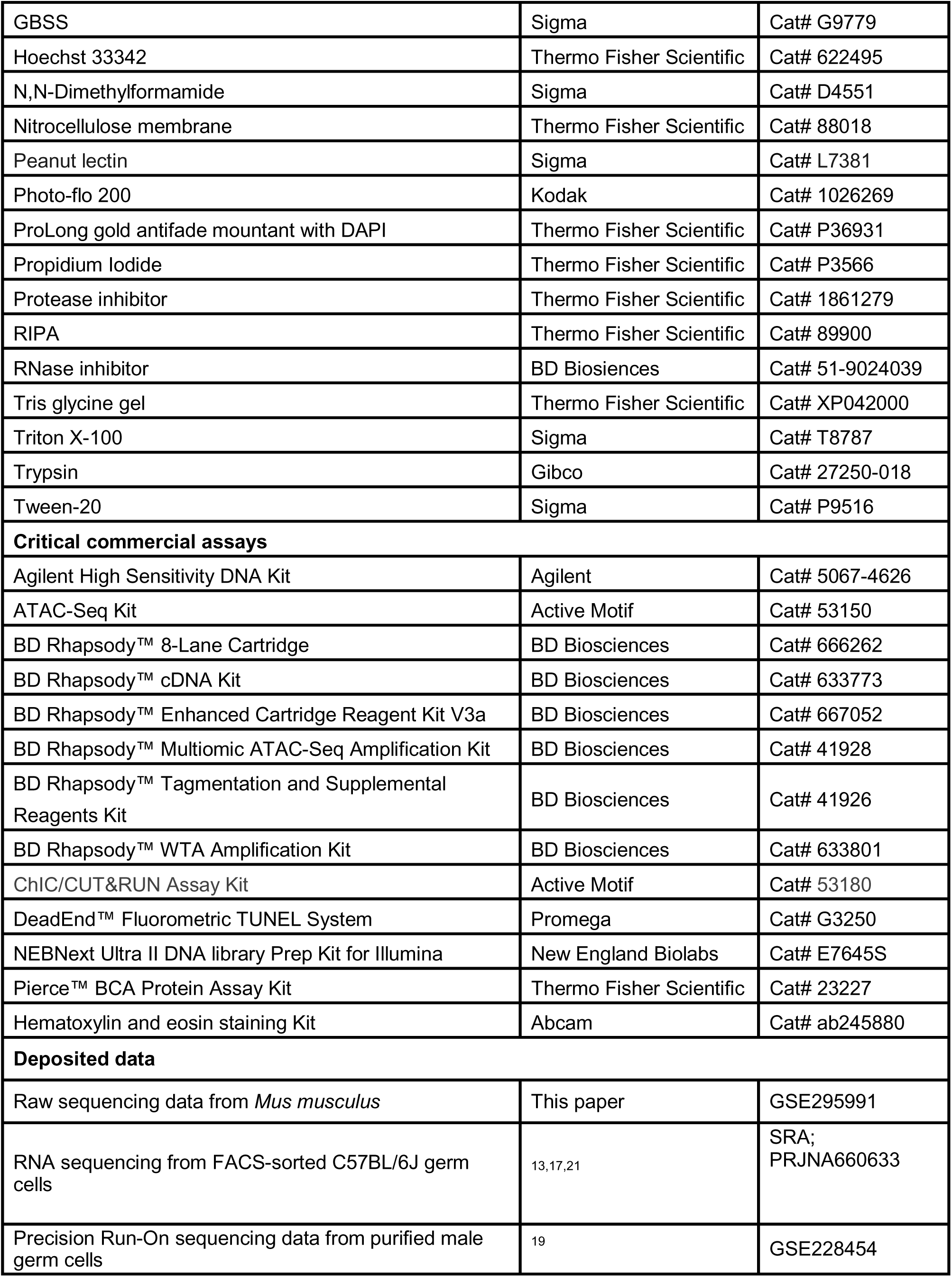
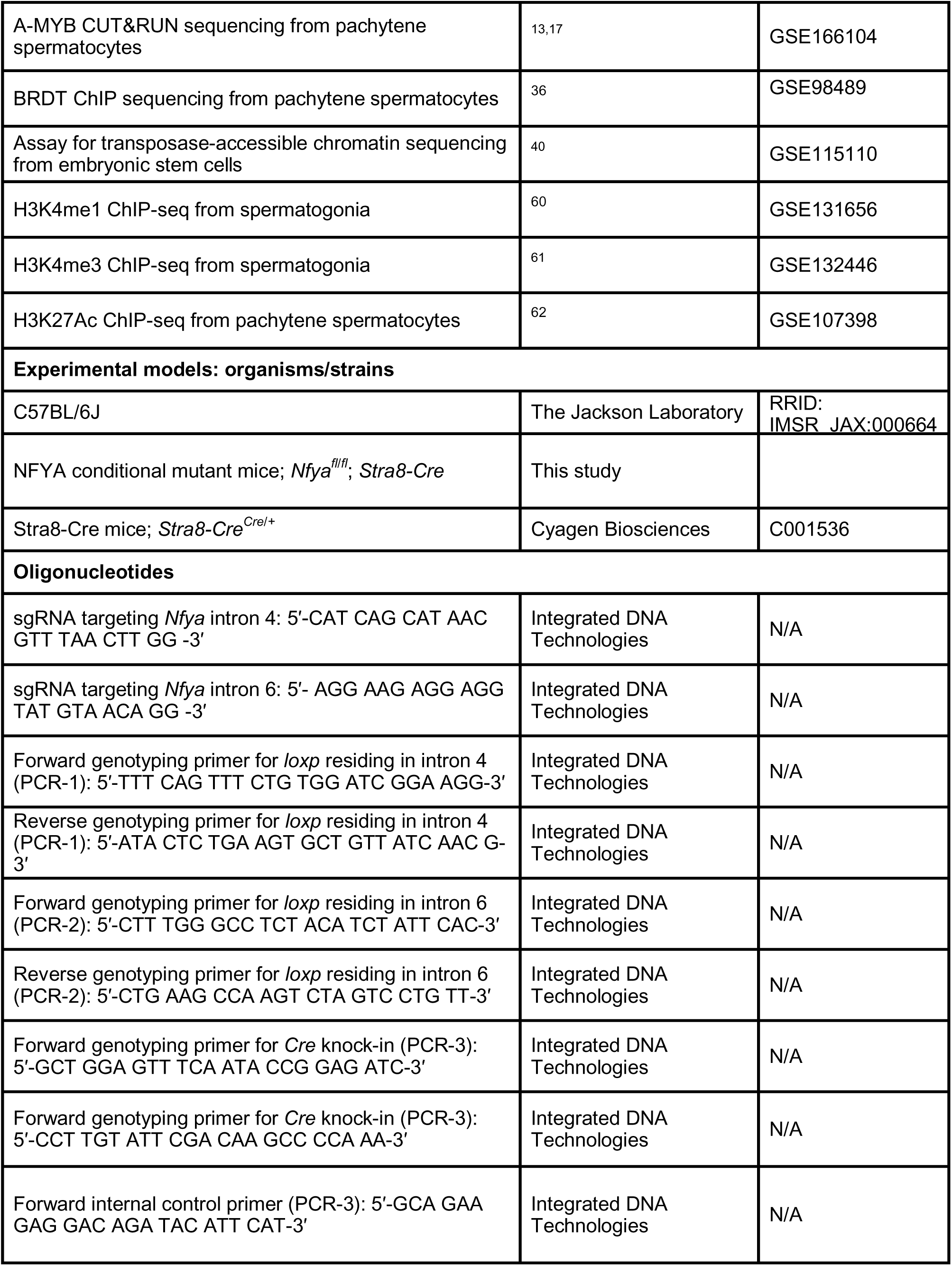
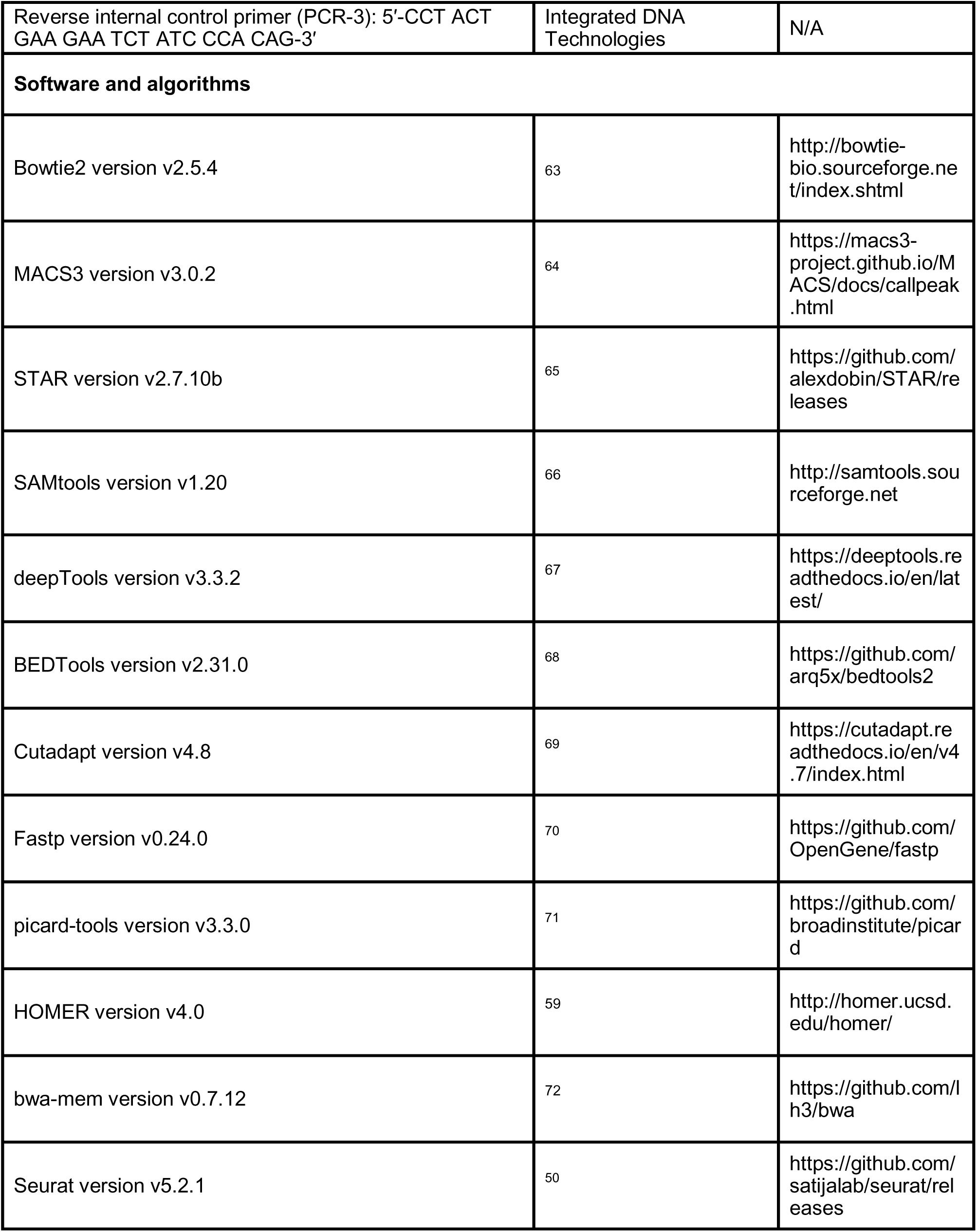
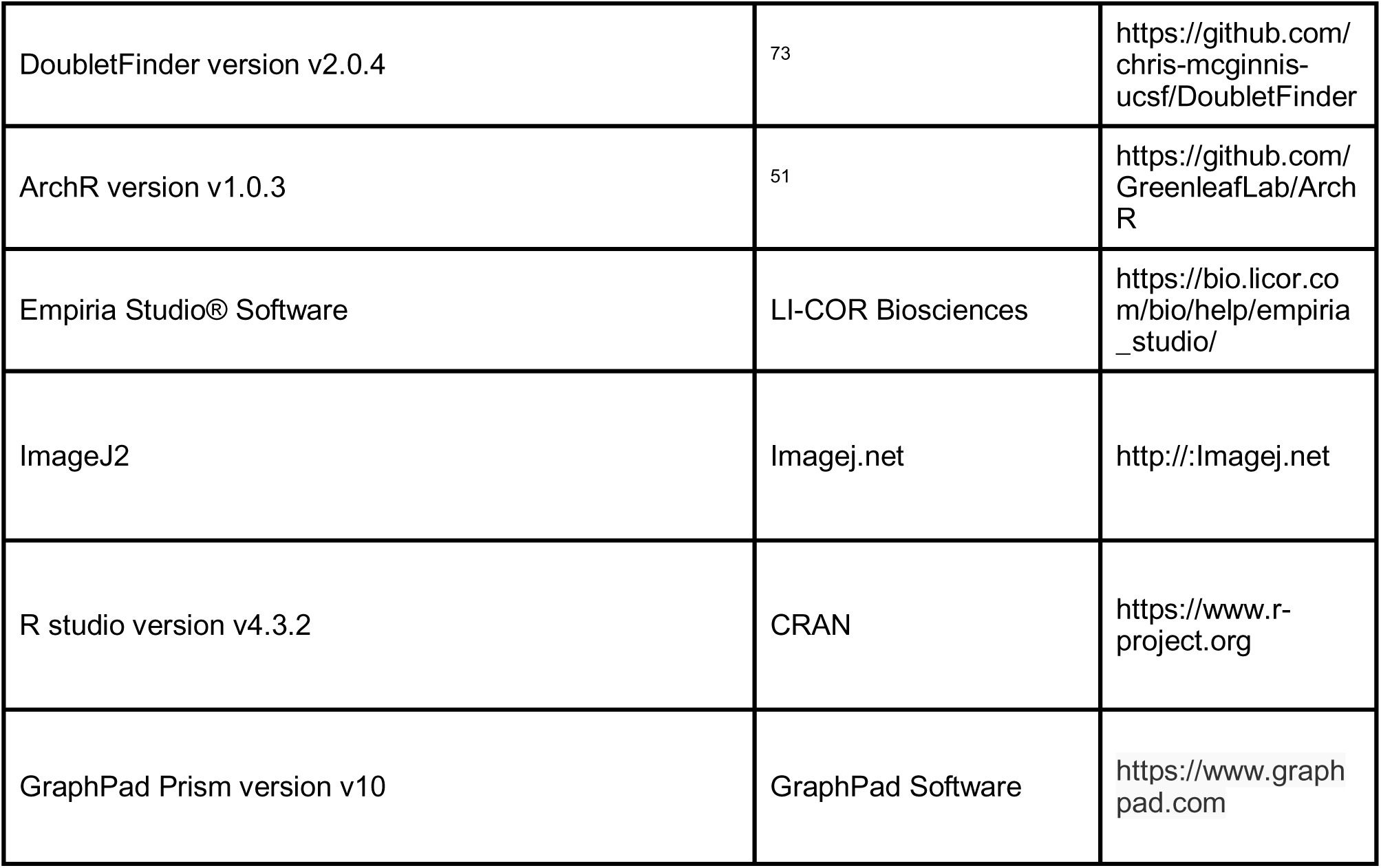

### EXPERIMENTAL MODEL AND SUBJECT DETAILS

#### Mice

Mice were maintained and used according to the guidelines of the Regional Animal Experimentation Ethics Committee of Swedish Board of Agriculture (8246-2021). C57BL/6J mice (RRID: IMSR_JAX:000664) were used as wild-type mice.

Male germ cell-specific *Nfya* mutant mice were generated by conditional deletion of exons 5–6 of *Nfya* using Stra8-Cre-loxP system (Cyagen Biosciences, Jiangsu, China). 5′ *loxP* was inserted in intron 4 and 3′ *loxP* was inserted in intron 6. Briefly, two guide RNAs targeting sequences in *Nfya* intron 4 (5′-CAT CAG CAT AAC GTT TAA CTT GG-3′) and intron 6 (5′-AGG AAG AGG AGG TAT GTA ACA GG-3′), the donor vector containing *loxP* sites, and Cas9 protein were co-injected into the cytoplasm of pronuclear stage embryos. The injected embryos were cultured in KSOM medium overnight and those which developed into the two-cell stage were transferred into the uterus of pseudo-pregnant ICR females. F0 founder mice were back-crossed with C57BL/6J mice in order to identify animals with germ line transmission in F1.

### METHOD DETAILS

#### Isolation of mouse germ cells by Fluorescence-Activated Cell Sorting

Male germ cell sorting was performed as previously described^13,17^ using the BD FACSMelody Cell Sorter (BD Biosciences). After dissecting testis from one mouse, the tubules were squeezed into 6 ml 1× Gey′s Balanced Salt Solution (GBSS, Sigma; G9779) containing 0.4 mg / ml collagenase type IV (Worthington; LS004188), which was then incubated at 33°C for 15 min at 600 rpm. Afterwards, the tubules were washed twice with 6 ml GBSS and incubated with 1× GBSS containing 0.5 mg/ml trypsin (Gibco; 27250-018) and 1 µg/ml DNase I (Roche; 10104159001) at 33°C for 15 min at 600 rpm. The tubules were then gently homogenized at 4°C on ice using Pasteur pipette. 400 μl FBS added to inactivate trypsin and the cells were passed through a pre-wetted 70 μm cell strainer (Corning; CLS431751). Cells were then pelleted in swing bucket centrifuge at 500 × g for 10 min at 4°C. Thereafter, cells were resuspended in 1× GBSS containing 5% (v/v) FBS, 1 µg/ml DNase I, and 5 μg/ml Hoechst 33342 (Thermo Fisher Scientific; 62249) and incubated at 33°C for 45 min at 150 rpm. The cell suspension was then placed on ice and propidium iodide (PI; 0.2 μg/ml, f.c.; Thermo Fisher Scientific; P3566) was added. Finally, cells were filtered through a pre-wetted 40 µm cell strainer (Corning; CLS431750). Four-way cell sorting was performed to obtain the populations of spermatogonia (SpG), leptotene and zygotene (L/Z), pachytene and diplotene (P/D), and round spermatids (RS) on BD FACSMelody Cell Sorter (BD Biosciences) using the gates as described in Figure S3A. 488-nm laser was used to excite PI, while 405-nm laser was used to excite Hoechst 33342. The emission for PI was recorded using 700/54 nm bandpass filter, while the emission of Hoechst 33342 was recorded by 660/10-nm (red) and 448/45-nm (blue) bandpass filters.

The purity of each sorted cell population was further assessed by chromosome spread followed by immunofluorescence staining on aliquots of cells. The purity of SpG population was assessed by DNA, GCNA, and DMRT1 staining, while the purity of L/Z, P/D, and RS was assessed by the pattern of DNA, γH2AX, and SYCP3 staining (Figure S3B). The purity of each population was calculated by assessing the staining from 42– 188 cells in six independent replicates (Figure S3B): SpG population, ∼95% spermatogonia; L/Z population, ∼5% spermatogonia, ∼50% leptotene spermatocytes, ∼29% zygotene spermatocytes, ∼10% early pachytene spermatocytes, <1% pachytene & diplotene spermatocytes and round spermatids, ∼5% spermatozoa; P/D population, <1% spermatogonia, ∼4% leptotene, zygotene, and early pachytene spermatocytes, ∼48% pachytene spermatocytes, ∼41% diplotene spermatocytes, ∼6% round spermatids, <1% spermatozoa R/S population, ∼88% round spermatids, ∼12% spermatozoa.

#### Chromosome spread

Cells were first incubated with 25 mM sucrose solution at room temperature for 20 min and then were fixed in 1% (w/v) paraformaldehyde solution containing 0.15% (v/v) Triton X-100 at room temperature for 2 h. We then added the cells to SuperFrost Plus microscope slides (VWR; 631-108) in a dropwise manner, in which the slides were pre-dipped in 1% (w/v) paraformaldehyde solution containing 0.15% (v/v) Triton X-100. Slides were then incubated for at least 2 h in a humidifying chamber. Afterwards, the lid of chamber was then removed to dry the slides completely. Dried slides were washed at room temperature for 10 min in a step-wise manner with (*i*) 1 × PBS containing 0.4% (v/v) Photo-Flo 200 (Kodak; 1026269), (*ii*) 1 × PBS containing 0.1% (v/v) Triton X-100, (*iii*) ADB/PBS solution [10% ADB (0.03% (w/v) BSA, 0.01% (v/v) Triton X-100, 10% (v/v) Donkey serum (Sigma, D9663), 2 × PBS) and 90% 1 × PBS). Slides were then incubated at 4°C overnight with primary antibodies diluted in ADB solution (anti-γH2AX, Novus Biologicals, NB100-384, 1:1000 dilution; anti-SCP-3 (D-1), Santa Cruz, sc-74569, 1:400 dilution; anti-GCNA, Abcam, ab82527 1:1000 dilution; anti-DMRT1, Santa Cruz, sc-377167, 1:200 dilution). Next day, slides were washed sequentially as (i–iii, above), and incubated at 37°C for 1 h with secondary antibodies diluted 1:2,000 in ADB (Alexa Fluor anti-mouse 594, Thermo Fischer Scientific , A21203; Alexa Fluor anti-rabbit 488, Thermo Fisher Scientific, A21206; Alexa Fluor anti-rat 594, Thermo Fisher Scientific, A21209). Finally, slides were washed three times in 1 × PBS containing 0.4% (v/v) Photo-Flo 200 at room temperature each for 10 min and once in 0.4% (v/v) Photo-Flo 200 at room temperature for 10 min. Slides were dried in dark and mounted with ProLong Gold Antifade Mountant with DAPI (Thermo Fisher Scientific, P36931). Images of chromosome spread were captured by Zeiss Axio Observer 7 inverted widefield microscope.

#### Testis histology by hematoxylin and eosin staining

Testis tissues from wildtype, *Cre/+,* and *Nfya-CKO* mice were fixed with Bouin’s solution (VWR; 7000.1000) at room temperature overnight. Next day, fixed tissues were washed with 70% (v/v) ethanol and embedded in paraffin to section them at 5 μm thickness. We performed hematoxylin and eosin (H&E) staining using H&E Staining Kit according to the manufacturer’s protocol (Abcam; ab245880). Images for H&E staining were captured using Zeiss Axio Observer 7 inverted widefield microscope.

#### Immunohistochemistry staining

Immunohistochemistry (IHC) was performed according to the standard protocols. In brief, we de-waxed 5 μm sections in xylene and dehydrated them in descending percentages of ethanol [100%, 95%, and 70% (v/v) ethanol; sections were incubated 5 min in each ethanol solution]. To retrieve antigens, sections were boiled in 1 mM citrate buffer (pH 6.0). The ImmPRESS Excel Amplified HRP Polymer Staining kit (Vector Laboratories; MP-7401) was used for IHC staining. To inactivate the endogenous peroxidase, tissue sections were incubated with 3% (v/v) hydrogen peroxide at room temperature for 10 min, then were blocked with 2.5% (v/v) horse serum. Sections were incubated with rabbit anti-NFYA (Sigma; HPA050779; 1:500 dilution) antibody at 4 °C overnight. Next day, sections were covered with secondary HRP anti-rabbit antibody (Vector Laboratories; MP-7401) and incubated at room temperature for 1 h. Following the secondary antibody incubation, chromogenic substrate (Thermo Fisher Scientific; TA-125-QHDX) was added and incubated at room temperature for 10 min. Thereafter, slides were counterstained with hematoxylin and dehydrated with ascending percentages of ethanol [70%, 95%, and 100% (v/v) ethanol]. Slides were finally covered with coverslips. IHC images were captured by Zeiss Axio Observer 7 inverted widefield microscope.

#### Immunofluorescence staining

##### Staining on Bouin’s fixed sections

5 μm tissue sections were deparaffinized in three sequential incubations in xylene for 7 min each and then washed in descending percentages of ethanol for 7 min each [100%, 95%, and 70% (v/v) ethanol]. Antigen retrieval was performed by boiling the slides in 1 mM citrate buffer (pH 6.0) at 95–100 °C for 20 min. We next cooled down the slides at room temperature for 10 min and washed them with 1 × PBS. Tissue sections on slides were outlined using PAP pen (Thermo Fischer Scientific; 008899). Following three washes in 1 × PBS (each wash was 5 min), sections were blocked with 5% (w/v) bovine serum albumin (Sigma; A9418) at room temperature for 1 h. Slides were then incubated at 4°C overnight with primary antibodies diluted in 1 × PBS (anti-DDX4/MVH, Abcam, ab13840, 1:500 dilution; anti-γH2AX, Novus Biologicals, NB100-384, 1:2000 dilution; FITC-conjugated peanut agglutinin, Sigma, L7381, 1:1000 dilution; anti-HSPA2, Sigma, A000798, 1:500 dilution; anti-SALL4, Abcam, ab57577, 1:1000 dilution; anti-c-KIT, Santa Cruz, sc-365504, 1:25 dilution; anti-NFYA (G-2), Santa Cruz, sc-17753 1:500 dilution; anti-STRA8, Proteintech, cL488-68071, 1:500 dilution). Following primary antibody incubation, slides were washed three times in 1 × PBS containing 0.02% (v/v) Tween-20 (Sigma; P9516) for 10 min each. Secondary antibodies were then applied at room temperature for 1 h (anti-rabbit Alexa Fluor 488, Sigma, A21206, 1:700 dilution; anti-rabbit Alexa Fluor 594, Sigma, A211012, 1:700 dilution; anti-mouse Alexa Fluor 594, Sigma, A21203, 1:700 dilution; anti-mouse Alexa Fluor 488, Sigma, A21202, 1:700 dilution). Nuclei were counterstained with DAPI (Sigma, D1306, 1:1000 dilution) at room temperature for 10 min. Finally, slides were washed twice with 1 × PBS for 10 min each and mounted with coverslips using DABCO mounting media (Sigma, D27802). Immunoflourescence images were captured by Zeiss Axio Observer 7 inverted widefield microscope.

##### Staining on cryosections

Testis tissues from *Cre/+,* and *Nfya-CKO* mice were fixed in 1 × PBS containing 4% (w/v) paraformaldehyde at 4°C overnight. Next day, fixed tissues were five times washed in 1 × PBS each for 30 min using a rocker platform. We then incubated the tissues in 30% (v/v) sucrose at 4°C overnight. Thereafter, tissues were embedded in FSC 22 Frozen Section Media (Leica, 3801480) and were frozen at -80°C. 5 μm cryosections were washed three times in 1 × PBS for 5 min each and proceeded with antigen retrieval as described for Bouin’s fixed sections. Slides were incubated with anti-SCP-3 (D-1) antibody (Santa Cruz; sc-74569; 1:2000 dilution) and the secondary antibody anti-mouse Alexa Fluor 594 (Sigma; A21203; 1:700 dilution). Both primary and secondary antibodies were diluted in 1 × PBS. Nuclei were counterstained with DAPI (Sigma, D1306, 1:1000 dilution) at room temperature for 10 min and slides were mounted with coverslips using DABCO mounting media (Sigma, D27802). Images for immunofluorescence staining on cryosections were captured by Zeiss Axio Observer 7 inverted widefield microscope.

#### TUNEL staining

The TdT-mediated dUTP Nick-End Labeling (TUNEL) assay was performed using the DeadEnd Fluorometric TUNEL System (Promega; G3250) according to the manufacturer’s protocol. 5 μm tissue sections were first deparaffinized in two sequential incubations in xylene for 5 min each and then washed in 100% (v/v) ethanol at room temperature for 5 min. We next rehydrated the sections by sequential incubation in descending percentages of ethanol for 3 min each [100%, 95%, 85%, 70%, and 50% (v/v) ethanol]. After rehydrating the samples, slides were washed once in 0.85% (w/v) NaCl and once in 1 × PBS at room temperature for 5 min each. The tissue sections were then fixed with 1 × PBS containing 4% (w/v) paraformaldehyde at room temperature for 15 min. After washing the slides two times in 1 × PBS for 5 min each, tissue sections were treated with 20 µg/ml Proteinase K at room temperature for 9 min. Following Proteinase K treatment, the tissue sections were washed once in 1 × PBS at room temperature for 5 min and fixed with 1 × PBS containing 4% (w/v) paraformaldehyde at room temperature for 5 min. Excess liquid was removed from the slides and tissue sections were covered with 100 μl Equilibration Buffer at room temperature for 5 min. Tissue sections were then incubated in 50 μl rTdT incubation buffer (45 μl Equilibration Buffer, 5 μl Nucleotide Mix containing fluorescein-12-dUTP, 1 μl rTdT Enzyme) in a humidifying chamber at 37°C for 1 h. To stop the reaction, the slides were immersed in 2 × saline-sodium citrate for 15 min followed by three sequential washes in 1 × PBS for 5 min each. Thereafter, tissue sections were stained with in 1 × PBS containing 1 μg/ml propidium iodide at room temperature for 15 min in the dark. Finally, slides were washed three times in deionized water for 5 min each and mounted with one drop of DABCO (Sigma; D27802), and the slides were covered with a glass coverslip. Images were captured using Zeiss Axio Observer 7 inverted widefield microscope.

#### Western blotting

Frozen tissues were finely minced and homogenized in a Dounce homogenizer using 30 strokes of pestle B in RIPA lysis buffer (Thermo Fisher Scientific; 89900; 25 mM Tris-HCl, pH 7.6, 150 mM NaCl, 1% (v/v) NP-40, 1% (w/v) sodium deoxycholate, and 0.1% (w/v) SDS) supplemented with 1% (v/v) protease inhibitor (Thermo Fisher Scientific; 1861279). The homogenized tissues were incubated on ice for 30 min and then centrifuged at 20,000 × *g* at 4°C for 30 min. The supernatant was collected into a new tube, and protein concentration was determined using the BCA Protein Assay Kit (Thermo Fisher Scientific; 23227). 75 µg total protein from each sample was mixed with ¼ volume of loading dye (Invitrogen; NP0007) containing 0.2M dithiothreitol and denatured at 95°C for 5 min. Proteins were resolved by electrophoresis through a 4– 20% Tris-Glycine gradient SDS gel (Thermo Fisher Scientific; XP04200) and transferred onto a 0.45 µm nitrocellulose membrane (Thermo Fisher Scientific; 88018). Membranes were blocked with Blocking Buffer (Thermo Fisher Scientific; 37565) at room temperature for 1.5 h. After blocking, the membranes were incubated at 4°C overnight with primary antibodies (rabbit anti-NFYA, Bethyl, A302-105A, 1:500; mouse anti-ACTIN, Sigma, A1978, 1:5000). Next day, membranes were washed three times with 1 × PBS-T (0.1% (v/v) Tween-20 in 1× PBS) for 30 min each, followed by incubation with secondary antibodies at room temperature for 1 h (anti-rabbit IRDye 800CW, LI-COR Biosciences, 926–32213, 1:15000; and anti-mouse IRDye 680RD, LI-COR Biosciences, 926–68072, 1:15000). After three additional washes with 1 × PBS-T for 15 min each, protein signal was detected using the Odyssey Infrared Imaging System (LI-COR).

#### ATAC-seq library construction

We performed <u>a</u>ssay for transposase-<u>a</u>ccessible <u>c</u>hromatin and <u>seq</u>uencing (ATAC-seq) according to the manufacturer’s protocol (Active Motif; 53150). Briefly, 100,000 FACS-purified spermatogonia (SpG), leptotene & zygotene (L/Z), and pachytene & diplotene (P/D) cells were pelleted by centrifuging at 500 × g at 4°C for 5 min. After carefully discarding supernatant, cells were gently washed with 100 μl ice-cold 1 × PBS which was followed by an additional centrifugation at 500 × g at 4°C for 5 min. After carefully removing 1 × PBS, cell pellet was resuspended in 100 μl ATAC lysis buffer which was immediately followed by centrifugation at 500 × g at 4°C for 10 min. After removing supernatant, nuclei were incubated in 50 μl Tagmentation Master Mix (25 μl 2 × tagmentation buffer, 12 μl water, 10 μl assembled transposase, 2 μl 10 × PBS, 0.5 μl 1% (v/v) Digitonin, and 0.5 μl 10% Tween-20) at 37°C for 30 min using PCR machine equipped with a heated lid. Immediately following transposition, tagmented DNA was mixed with 250 μl DNA binding buffer and 5 μl sodium acetate. The mixture was then added onto a column and centrifuged at 17,000 × g for 1 min. Column was then washed once with 750 μl wash buffer and the tagmented DNA was eluted from the column with DNA Purification Elution Buffer. After purifying tagmented DNA, the library of fragments was amplified in 1 × NEB Q5 PCR master mix (1 × Q5 Reaction buffer [NEB; B9027S], 1 μM Illumina i7 Indexed primer, 1 μM Illumina i5 Indexed primer, 0.2 mM dNTPs, and 0.5 U Q5 High-Fidelity DNA polymerase [NEB; M0491S]). The following PCR condition was applied to generate ATAC-seq libraries: 72°C for 5 min; 98°C for 30 s; 10 cycles of thermocycling at 98°C for 10 s, 63°C for 30 s, and 72°C for 1 min. ATAC-seq libraries were sequenced as 79 + 79 nt paired-end reads using NextSeq550 system (Illumina).

#### Simultaneous single-cell ATAC-seq and mRNA-seq profiling

We performed BD Rhapsody Single-Cell Analysis System to simultaneously analyze the profile of mRNA expression and accessible chromatin state in cells from the testis of 11, 17 and 25 dpp wild-type mice and 8-weeks-old *Cre/+* and *Nfya-CKO* mice.

##### Nuclei isolation for single-cell multiomics

Testis were dissected and squeezed into 6 ml 1× Gey′s Balanced Salt Solution (GBSS, Sigma; G9779) containing 0.4 mg/ml collagenase type IV (Worthington; LS004188) and incubated at 33°C for 15 min at 600 rpm. We then washed the tubules twice with 6 ml GBSS and incubated them in 1 × GBSS containing 0.5 mg/ml trypsin (Gibco; 27250-018) and 1 µg/ml DNase I (Roche; 10104159001) at 33°C for 15 min at 600 rpm. We thereafter homogenized the tubules using Pasteur pipette at 4°C on ice for 3 min. 400 μl FBS added to inactivate trypsin and the cells were passed through a pre-wetted 70 μm cell strainer (Corning; CLS431751). Cells were then pelleted in swing bucket centrifuge at 500 × g for 10 min at 4°C and resuspended in 3.5 ml 1 × GBSS, 175 μl FBS, 6 μl DNase I. After resuspension, the cells were filtered through a pre-wetted 40 μm cell strainer (Corning; 43750). 300,000 cells were used for nuclei isolation. Cells were pelleted at 400 × g for 10 min at 4°C and gently resuspended in 195 μl swelling buffer (10 mM (w/v) Tris-HCl pH 7.5, 2 mM (w/v) MgCl2, 3 mM (w/v) CaCl2). Additional 1.8 ml swelling buffer was added to resuspended cells dropwise. Cells were then incubated on ice for 5 min and pelleted at 400 × g for 10 min at 4°C. Supernatant was carefully removed and cell pellet was resuspended in 100.5 μl prechilled lysis buffer *1* (9 mM (w/v) Tris-HCl pH 7.5, 1.8 mM (w/v) MgCl2, 2.7 mM (w/v) CaCl2, 10% (v/v) glycerol, 1 × protease inhibitor cocktail (PIC), 0.4 U/μl RNasin [Promega; N2615]). Afterwards, 100.5 μl lysis buffer *2* (9 mM (w/v) Tris-HCl pH 7.5, 1.8 mM (w/v) MgCl2, 2.7 mM (w/v) CaCl2, 10% (v/v) glycerol, 10% (v/v) Igepal CA-360, 1 × PIC, 0.4 U/μl RNasin [Promega; N2615]) was added dropwise to the mix and incubated on ice for 5 min. The nuclei were pelleted at 600 × g at 4°C for 5 min. The supernatant was discarded, and the nuclei were resuspended in 200.25 μl lysis buffer *3* (9 mM (w/v) Tris-HCl pH 7.5, 1.8 mM (w/v) MgCl2, 2.7 mM (w/v) CaCl2, 10% (v/v) glycerol, 0.01% (v/v) Digitonin, 1 × PIC, 0.4 U/μl RNasin [Promega; N2615]). Afterwards, additional 1.8 ml lysis buffer *3* was added dropwise and carefully mixed. The nuclei were centrifuged at 500 × g at 4°C for 5 min. The supernatant was removed, and the nuclei were resuspended in 200.25 μl freezing buffer (50 mM (w/v) Tris-HCl pH 8, 5 mM (w/v) MgCl2, 40% (v/v) glycerol, 1 × PIC, 0.4 U/μl RNasin [Promega; N2615]). The nuclei were centrifuged at 4°C at 900 × g for 6 min. The supernatant was discarded, and the nuclei were resuspended in 50 μl modified nuclei buffer (BD Biosciences; 51-9023091) supplemented with 2.5% (v/v) RNase inhibitor (BD Biosciences; 51-9024039) and 1 mM (v/v) DTT. ATAC-seq and mRNA-seq libraries were prepared from 50,000 nuclei.

#### Simultaneous ATAC-seq and mRNA whole transcriptome (WTA) library preparation

To tagment DNA, 50,000 nuclei in 5 μl modified nuclei buffer (BD Biosciences; 667052) were mixed with 11.75 μl water, 2 μl 10 × PBS, 1.25 μl RNase inhibitor (BD Biosciences; 51-9024039), 0.5 μl 1% (v/v) Digitonin, 0.5 μl 10% (v/v) Tween-20, 4 μl Tagmentase, and incubated at 37°C for 30 min. After incubation, 300 μl modified sample buffer was added to the mix and the nuclei were loaded into the BD Rhapsody 8-Lane Cartridge (BD Biosciences; 666262). The cartridge was incubated at room temperature for 8 min to allow nuclei to sediment in the micro-wells of cartridge. Nuclei were then washed twice with cold sample buffer and incubated with splint beads (BD Biosciences; 667052) at room temperature for 3 min. Next, the cartridge was quickly agitated at room temperature for 10 sec at 1000 rpm using ThermoMixer (Eppendorf; EP5382000015) and washed twice with cold sample buffer. Afterwards, 280 μl lysis buffer, supplemented with 10 mM DTT and 25 μl proteinase K, was added and incubated at 37°C 10 min. After the lysis, the tagmented DNA and mRNA were immobilized onto beads via splint-oligo-bonded TSO and poly(T) oligo, respectively. We thereafter collected the beads retaining tagmented DNA and mRNA into a 1.7 ml tube, where we first ligated tagmented DNA to BD Rhapsody bead oligo in ligation mix (20 μl 10 × ligation buffer, 10 μl DNA ligase [BD Biosciences; 41928], 5 μl RNase inhibitor [BD Biosciences; 51-9024039], 170 μl water) at 25°C for 30 min at 1200 rpm. Afterwards, the ligation mix was discarded and reverse transcription (RT) was performed in 200 μl RT mix (40 μl RT buffer [BD Biosciences; 633773], 20 μl 10 mM dNTPs, 10 μl 0.1M DTT, 12 μl RT/PCR enhancer, 10 μl RNase inhibitor [BD Biosciences; 633773], 10 μl reverse transcriptase [BD Biosciences; 633773], 98 μl water) was incubated at 42°C for 30 min at 1200 rpm. Following RT reaction, RT mix was replaced by splint-oligo removal buffer and the tube was incubated at 60°C for 5 min at 1200 rpm. To remove unused oligos on the beads, samples were then incubated in 200 μl Exonuclease I mix (20 μl 10 × exonuclease I buffer, 10 μl Exonuclease I [BD Biosciences; 633773], 170 μl water) at 37°C for 30 min at 1200 rpm. We then quenched Exonuclease I by adding 4 μl 0.5 M EDTA to mix. To prepare ATAC-seq libraries, two rounds of DNA elution (80 μl elution) from the beads was performed. Note that after eluting DNA, beads were stored in bead resuspension buffer at 4°C to later prepare WTA libraries. 80 μl ATAC-seq libraries were amplified and indexed in 42 μl PCR index mix (30 μl PCR master mix [BD Biosciences; 41928], 6 μl ATAC-seq forward primer, 6 μl ATAC-seq reverse primer) with the following PCR condition: 98°C 45 s, 12 cycles of thermocycling at 98°C 10 s, 66°C 30 s, 72°C 30 s, and a final extension at 72°C for 1min.

To prepare WTA libraries, the beads, which were stored at 4°C, were first heated at 95°C for 5 min. Using magnet, the supernatant was discarded and beads were incubated with 87 μl Random Primer mix (10 μl WTA extension buffer [BD Biosciences; 633801], 10 μl WTA extension primer, 67 μl water) at 95°C for 5 min, followed by sequential incubation at 37°C and 25°C for 5 min each at 1200 rpm. Next, 13 μl Extension Enzyme mix (8 μl 10 mM dNTP, 12 μl ET/PCR enhancer, 6 μl WTA Extension Enzyme [BD Biosciences; 633801]) was added, and the mixture was incubated in ThermoMixer (Eppendorf; EP5382000015) at 1200 rpm with the following program: 25°C 10 min, 37°C 15 min, 45°C 10 min, 55°C 10 min. We then incubated the beads in 205 μl elution buffer at 95°C for 5 min to purify Random Primer Extension (RPE) product from the beads. Note that WTA Random Priming and Extension, followed by purification, was repeated once more. Total of 400 μl RPE product was purified with 1.8 × AMPure XP Beads (Beckman Coulter; A63881) and eluted in 40 μl Elution buffer. After elution, we added 80 μl PCR Master Mix containing 10 μl Universal oligo and 10 μl WTA Amplification primer directly to 40 μl eluted RPE product to perform initial amplification of WTA libraries with the PCR condition: 95°C 3 min, 11 cycles of thermocycling at 95°C 30 s, 60°C 1 min, 72°C 1 min and a final extension at 72°C for 2 min. The reaction was cleaned using 0.8 × AMPure XP Beads (Beckman Coulter; A63881). Before performing the final amplification, WTA libraries were analyzed on Bioanalyzer (Agilent; 2100 Bioanalyzer) to calculate the molar concentration of the DNA ranging from 150 to 600 bp. 10 μl 2 nM WTA libraries were indexed in 40 μl PCR index mix (25 μl PCR master mix [BD Biosciences; 633801], 5 μl BD Rhapsody library forward primer, 5 μl BD Rhapsody library reversed indexed primer, 5 μl water) with following PCR condition: 95°C 3 min, 8 cycles of thermocycling at 95°C 30 s, 60°C 30 s, 72°C 30 s, and a final extension at 72°C for 1 min.

ATAC-seq and WTA libraries were sequenced using NovaSeq X Plus Platform (Illumina) with the following sequencing setup: for ATAC-seq libraries, Read 1, 50 cycles, Read 2, 50 cycles, Index 1, 8 cycles, Index 2, 60 cycles; for WTA libraries, Read 1, 51 cycles, Read 2, 71 cycles, Index 1, 8 cycles. We sequenced the libraries with the following manufacturer’s recommended sequencing depth: for ATAC-seq libraries, 50,000 read pairs per cell; for WTA libraries, 100,000 read pairs per cell. Sequencing statistics are provided in Table S5.

#### CUT&RUN Sequencing

Cleavage Under Targets and Release Using Nuclease (CUT&RUN) sequencing from FACS-purified germ cells was performed according to the manufacturer’s protocol (Active Motif; 53180). 500,000 FACS-purified spermatogonia (SpG), leptotene & zygotene (L/Z), and pachytene & diplotene (P/D) cells were pelleted by centrifugation at room temperature for 3 min at 600 × g. The cell pellet was then gently resuspended in 100 μl nuclear isolation buffer (20 mM (w/v) HEPES-KOH, pH 7.9, 10 mM (w/v) KCl, 0.5 mM (v/v) Spermidine, 0.1% (v/v) Triton X-100, 20% (v/v) glycerol) supplemented with 1 × protease inhibitors, 0.5 mM Spermidine) and incubated on ice for 10 min. After incubation, cells were centrifuged at 600 × g at 4°C for 3 min. After removing the supernatant carefully, nuclei were washed once and resuspended in 100 μl wash buffer. 100 μl nuclei suspension was incubated at room temperature for 10 min with 10 μl Concanavalin A-coated paramagnetic beads to immobilize nuclei on beads. After immobilizing beads, supernatant was removed using magnet and 50 μl antibody buffer containing 1 μg anti-NFYA (Bethyl; A302-105A) or 0.5 μg IgG (EpiCypher; 13-0042) antibodies were added directly to beads. Samples were incubated at 4°C overnight on a nutator. Next day, beads were washed twice with 200 μl cell permeabilization buffer (CPB) and resuspended in 50 μl CPB. Afterwards, 2.5 μl ChIC/CUT&RUN pAG-MNase was added to each sample, and samples were incubated at room temperature for 10 min. Beads were then washed twice with 200 μl CPB and resuspended in 50 μl CPB. We then added 1 μl 0.1 M CaCl_2_ was added to each sample and incubated them at 4°C for 2 hours on a nutator. The reaction was stopped by adding 40 μl STOP solution and incubated at 37°C for 10 min in a thermal cycler whose lid was at 65°C. Samples were then placed on a magnet and the clear supernatant containing the DNA fragments were moved to a new tube. DNA was then purified by column purification provided with the kit. After purifying the DNA, CUT&RUN libraries were prepared using NEBNext Ultra II DNA Library Prep Kit (NEB; E7645) according to the manufacturer’s protocol. CUT&RUN libraries were sequenced as 79 + 79 nt paired-end reads using NextSeq550 system (Illumina).

#### Long RNA library analysis

RNA-seq analysis was performed as previously described^13,17,74^. Molecules of transcripts per cell was calculated as previously described^74^. Given that each sample contained ∼623,291,645 molecules of ERCC spike-in mix, the abundance of each gene = (number of mapped reads × 623291645)/(number of cells used to prepare the library × the number of reads mapping to the ERCC spike-in sequences).

#### PRO-seq analysis

We re-analyzed publicly available, GSE228454, <u>P</u>recision <u>R</u>un-<u>O</u>n sequencing (PRO-seq) data from purified spermatogonia (SpG), primary spermatocytes (SpI), and round spermatids (RS). Adapter sequences were trimmed from raw read pairs using Cutadapt (v4.8^69^) with a 10% error rate threshold. Afterwards, additional one nucleotide was removed from the end of read 1 (R1). Subsequently, reads were aligned to mouse reference genome (mm10) and *drosophila melanogaster* reference genome (dm6), which served as spike-in control, using Bowtie2 (v2.5.4^63^) with the parameters `—local —very-sensitive —no-unal —no-mixed —no-discordant -|10 -X700`. We specified insert size ranging from 10 to 700 bp to ensure accurate and high-confidence alignments. SAM files were converted to BAM format using SAMtools (v1.20^66^), and BAM files were subsequently sorted, indexed, and filtered to retain only uniquely mapped reads. Final indexed BAM files were then converted into bigwig format using deepTools (v3.3.2^67^). Each bigwig file was normalized to scale factor. Scale factor was calculated by dividing the number of spike-in reads by the total number of mapped reads. Because round spermatids have haploid genome, scale factor for RS was halved. Metagene plots were generated using PRO-seq reads normalized to scale factor with `plotProfile` function from deepTools (v3.3.2^67^). PRO-seq statistics are provided in Table S5.

To examine the PRO-seq peaks near transcription start sites (TSSs) of genes, peaks were called using MACS3 (v3.0.2^64^; FDR < 0.1) with parameters `—q 0.1 —min-length 150` in each biological replicates from SpG, SpI, and RS. Bed file containing genomic coordinates for all annotated TSSs was retrieved from UCSC Genome Browser (mm10; GENCODE VM23). Distance from the TSS to the nearest PRO-seq peak summit for each gene was calculated. If a meiosis-I or spermiogenesis gene had a PRO-seq peak ±2 kb of its TSS in at least two replicates of PRO-seq from SpG, it was considered as *poised* gene. Note that for those genes with multiple TSSs, poised classification was assigned if at least one of the TSSs had a PRO-seq peak within ±2 kb in at least two biological replicates.

To study promoter-proximal RNA polymerase II (Pol II) pausing, we quantified PRO-seq reads at promoter-proximal regions and gene bodies of genes using BEDTools (v.2.31.0^68^). Promoter-proximal region is defined as the first 5% of the gene from its transcription start site, whereas gene body is defined as a region starting from the first 30% a gene to the transcription end site (TES). The pausing index is calculated by dividing the reads at promoter-proximal region by gene body reads. Two-sided wilcoxon matched-pairs signed ranked sum test was applied to measure the statistical difference for the same gene categories between different germ cells.

#### ATAC-seq analysis

Adapter sequences and low-quality reads were removed from the raw ATAC-seq reads using Fastp (v. 0.24.0^70^). Subsequently, reads were aligned to the mouse reference genome (mm10) using Bowtie2 (v2.5.4^63^) with the parameters `—very-sensitive —no —unal —no-mixed —no-discordant`. Multiple aligning read pairs were removed using SAMtools (v1.20^66^). We then identified and removed duplicate reads using `MarkDuplicates` function from picard-tools (v.3.3.0^71^). BAM alignments were then converted to bigwig format using deepTools (v.3.3.2^67^) and normalized to reads per million mapped reads (RPM) accounting for differences in sequencing depth. ATAC-seq peaks were called in each biological replicates from spermatogonia (SpG), leptotene/zygotene (L/Z), and pachytene/diplotene (P/D) using MACS3 (v3.0.2^64^; FDR < 0.01). ATAC-seq statistics are provided in Table S5.

Distance from the TSS to the nearest ATAC-seq peak summit for each gene was calculated. Genes that retain ATAC-seq peak ±2 kb of their TSSs in at least two replicates of ATAC-seq experiments were considered as genes with accessible chromatin. HOMER (v4.0^59^) was employed to identify transcription factor binding motifs under ATAC-seq peaks around the promoters of poised genes in SpG cells. Motif analysis scanning ±75 bp from the center of each peak was allowed for up to 25 motifs per peak. Promoters of genes, whose transcript abundance remained constant across germ cells, served as background control (Table S1).

#### CUT&RUN analysis

Adapter sequences were trimmed and low-quality reads were removed from the raw paired-end CUT&RUN reads using Fastp (v0.24.0^70^). Reads were then aligned to the mouse reference genome (mm10) using Bowtie2 (v2.5.4^63^) with parameters `—local — very-sensitive —no-unal —no-mixed —no-discordant -|10 -X700`. Subsequently, SAM files were converted to BAM format using SAMtools (v1.20^66^), and alignments whose quality is less than 20 were removed. BAM files were then sorted and indexed, and multiple aligning read pairs were removed. Thereafter, duplicate reads were removed using `MarkDuplicates` function from picard-tools (v.3.3.0^71^). Bigwig files were generated using `bamCoverage` function from deepTools (v3.3.2^67^) and normalized to reads per million (RPM). CUT&RUN statistics are provided in Table S5.

NFYA peaks were identified using MACS3 (v3.0.2^64^; FDR < 0.1). CUT&RUN for IgG antibody was used as the background control to call significant NFYA peaks. Genes with NFYA peak within ±2 kb of their transcription start sites in at least two replicates of CUT&RUN experiments were considered as NFYA-bound genes.

#### Simultaneous single-cell ATAC-seq and mRNA-seq analysis

##### Cell barcode annotation

Valid cell barcodes were downloaded from BD Biosciences. For scRNA-seq, read 1 (R1) contains the information of barcodes and unique molecular indices (UMIs), while read 1 (R2) are from complementary DNA (cDNA). R1 retains 35 to 38 nucleotides cell barcodes followed by 8 nucleotide UMIs. For scATAC-seq, read I1 (Rl1) marked barcode and UMIs, while R1 and R2 are paired DNA sequences. The construction of Rl1 is 35 nucleotides cell barcodes followed by 8 nucleotides UMIs in reverse complemented. Allowing 0 mismatch, raw reads without valid cell barcodes were filtered. For the remaining reads, information of cell barcodes and UMIs were assigned to the read names of the cDNA end (scRNA-seq) or the paired DNA end (scATAC-seq).

##### Read alignment

We used STAR (v2.7.10b^65^) to align scRNA-seq reads to the mouse reference genome (mm10) using parameters `--outFilterIntronMotifs RemoveNoncanonicalUnannotated –outFilterMultimapNmax 1`. PCR duplications with the same UMIs and same alignment coordinates from the same cells were removed. Afterwards, a cell-by-gene sparse matrix was made for each sample. We used bwa-mem (v0.7.12^72^) to align scATAC-seq reads to the mouse reference genome (mm10) using default parameters. The aligned files were converted to bed format using BEDTools (v2.27.1^68^), with +4/–5 coordinate shift for plus/minus strand reads to account for the 9 bp apart cuts by Tn5 transposase. PCR duplications with the same UMIs and same alignment coordinates from the same cells are removed. A fragment file was then generated for each sample.

##### Cell type annotation

Cell type annotation was done using the Seurat (v5.2.1^50^) R package for scRNA-seq. We first removed cells with <500 expressed genes or ≥5% UMIs from mitochondrial genes. The gene UMI counts for each of the remaining cell were log normalized with a factor equals to 10,000. Top 5,000 variable genes were then extracted for principal component analysis (PCA). Top 20 PCs were further used for clustering (resolution = 0.8) and uniform manifold approximation and projection (UMAP) analysis. For each sample, we then used the DoubletFinder R package (v2.0.4^73^) to remove doublets and multiplets using the top 10 PCs and default parameters. A total of 33 cell clusters were identified. We assigned cell-type to each cell cluster using the expression of marker genes (Figure S9B; Table S4B). Clusters without clear marker gene expression were removed from downstream analysis.

Cell-type annotation for scATAC-seq was performed using ArchR R package (v1.0.3^51^). We filtered cells with <1,000 fragments or a TSS score <4. Doublets/multiplets were removed using ArchR for each sample with default parameters. Genome-wide tile matrix of 500 bp and UMAP analysis with default parameters were used for clustering. A total of 25 cell clusters were identified. We then assigned cell-type for each cluster based on the cell-type annotation generated from scRNA-seq analysis for those cells with both open chromatin and transcriptome captured along with the chromatin accessibility of marker genes.

##### Pseudo-bulk analysis

According to the cell-type annotation, we generated pseudo-bulks by merging all aligned reads/fragments for each cell type for scRNA-seq and scATAC-seq, respectively. We then normalized each pseudo-bulks to reads/fragments per million mapped reads/fragments and generated bigwig tracks. Chromatin accessibility of gene promoters is defined by the average normalized coverage at ±200 bp from TSS.

#### ChIP-seq analysis

Re-analysis and peak calling for publicly available A-MYB CUT&RUN (GSE166104^13,17^) and BRDT Chromatin Immunoprecipitation and sequencing (ChIP-seq) (GSE98489^36^) were performed by the pipeline used for CUT&RUN. A-MYB and BRDT signal intensities were quantified using BEDTools (v2.31.0^68^) within a ±2 kb of the transcription start sites (TSSs) of meiosis-I and spermiogenesis genes. Violin plots were generated using R (v4.3.2) to visualize the distribution of signal intensities at the promoters of poised and non-poised meiosis-I and spermiogenesis genes. Mann-Whitney U test was used to measure the significance for the differences between poised and non-poised genes in respect to A-MYB or BRDT occupancy.

To identify distal enhancer regions in the genomes of spermatogonia (SpG) and pachytene/diplotene spermatocytes (P/D), we re-analyzed publicly available H3K4me1 (GSE131656^60^) and H3K4me3 (GSE132446^61^) ChIP-seq data from SpG, as well as H3K27Ac (GSE107398^62^) ChIP-seq data from P/D, along with our H3K4me3 CUT&RUN from P/D. ChIP-seq data processing was performed according to the pipeline used for CUT&RUN analysis. Because H3K4me3 is associated with promoter-proximal regions, while H3K4me1 and H3K27Ac are found both at distal enhancers and promoter-proximal regions^75,76^, we employed exclusion criterion to identify distal enhancer regions. Specifically, for SpG cells, we excluded H3K4me1 peaks that overlapped with H3K4me3 peaks. Similarly, for P/D cells, H3K27Ac peaks overlapping with H3K4me3 were removed. Remaining H3K4me1 and H3K27Ac were then considered as distal enhancer regions. Subsequently, those NFYA peak summits that resides within the coordinates of H3K4me1 and H3K27Ac peaks non-overlapping with H3K4me3 were considered as NFYA peaks at enhancers.

### QUANTIFICATION AND STATISTICAL ANALYSIS

Statistical analyses and graph generation were conducted using R v4.3.2 (https://www.rstudio.com) or Prism 10.1.1 (GraphPad Software, LLC). Box plots were used to present data distribution. Boxes represent interquartile ranges (IQRs), spanning the first and third quartiles. Outliers were defined as data points with values > third quartile + 1.5 × IQR or lower than the first quartile − 1.5 × IQR, where IQR is the difference between the maximum of the third and the minimum of the first quartile. Relationship between variables were measured using spearman’s correlation (p). Two-sided wilcoxon matched-pairs signed ranked sum test was used to calculate *p* values in Figures 1D, 6D, 7C and S6E. Two-sided Mann-Whitney-Wilcoxon U test was used to calculate *p* values for Figure S2D. Two-sided unpaired t test was used to calculate *p* values in Figures 4B-4G, S8B, S8D-S8F.

