## Supplementary figures and images for "Transcription factor NFYA directs male meiotic entry by facilitating accessible chromatin at meiotic promoters in mice"

### Figure S1

Säflund et al., Figure S1

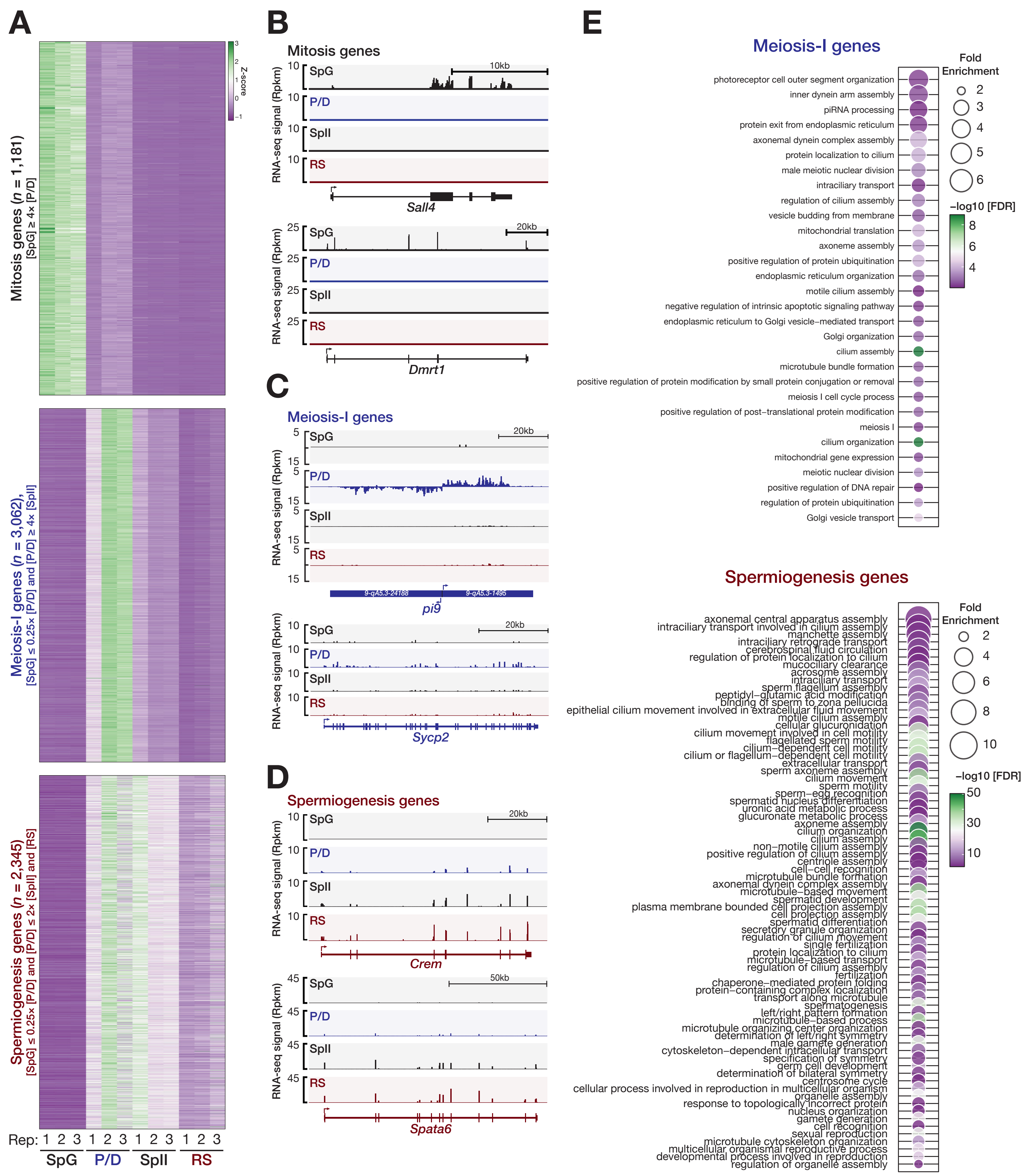

### Figure S2

Säflund et al., Figure S2

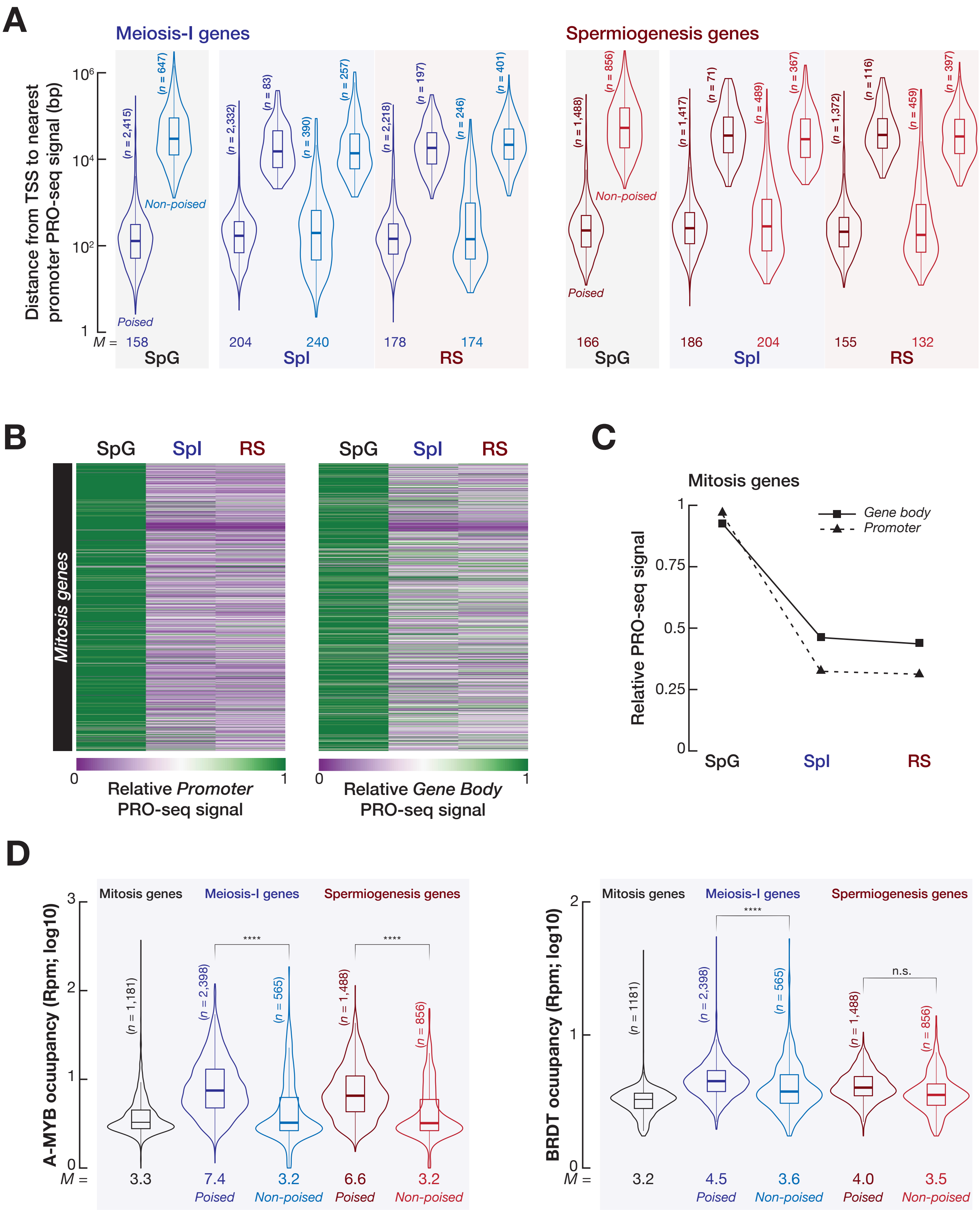

### Figure S4

Säflund et al., Figure S4

A

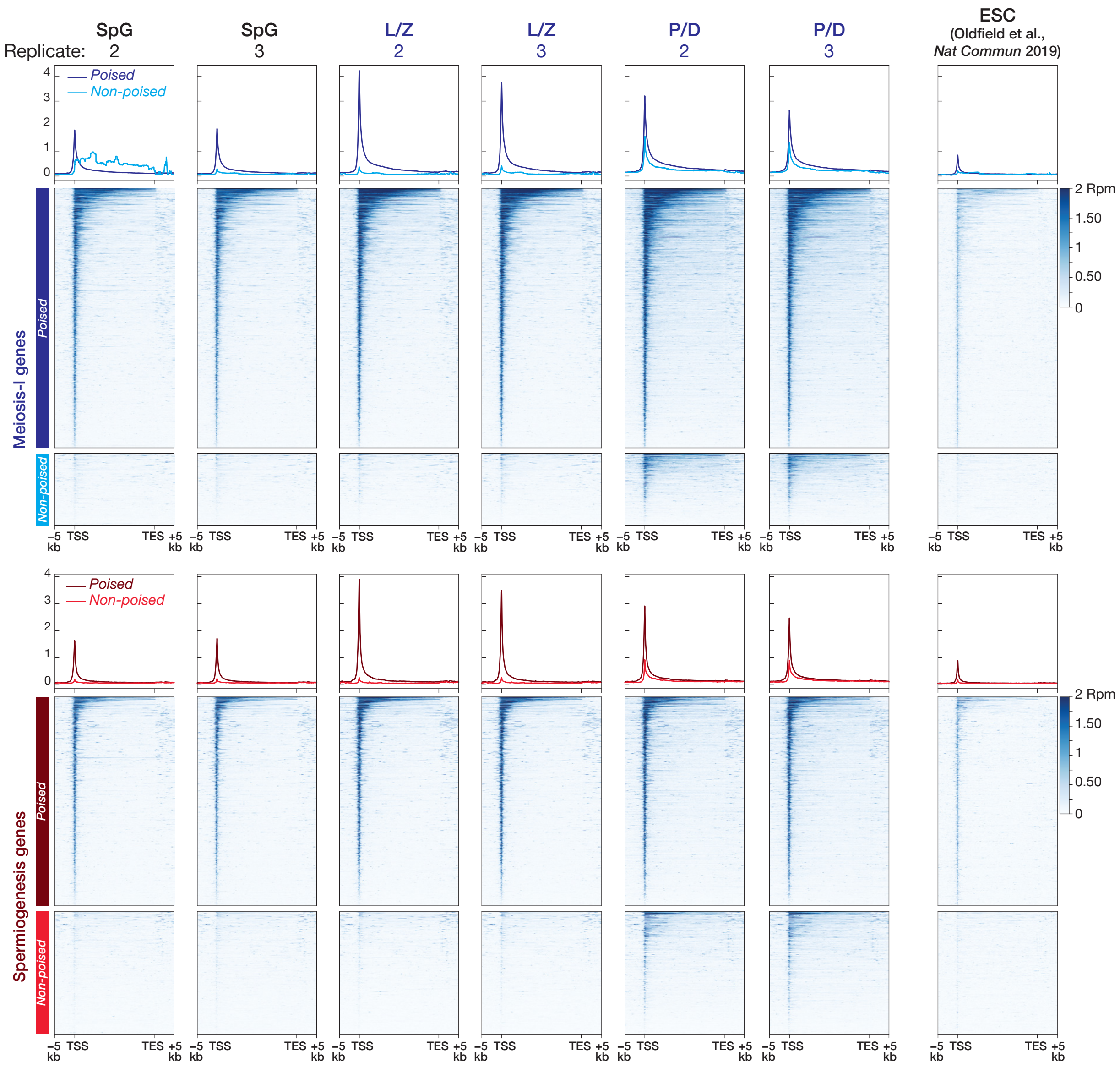


B

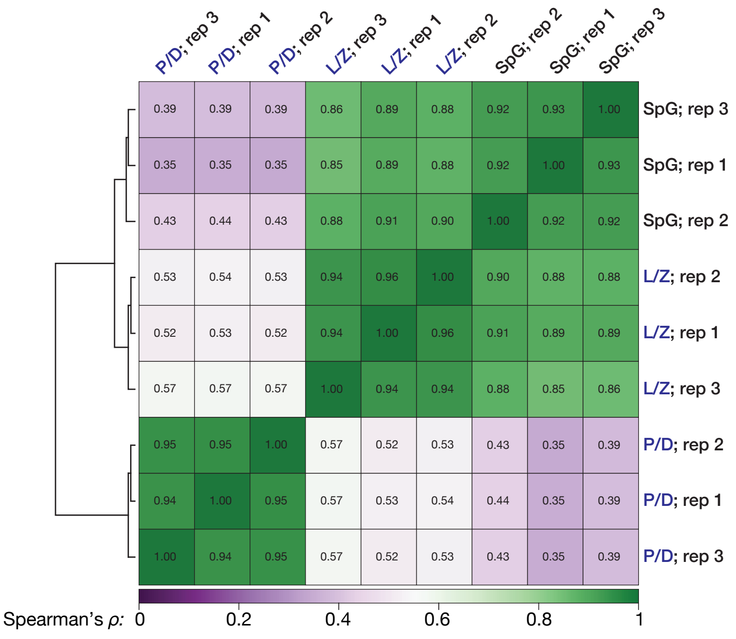

### Figure S5

Säflund et al., Figure S5

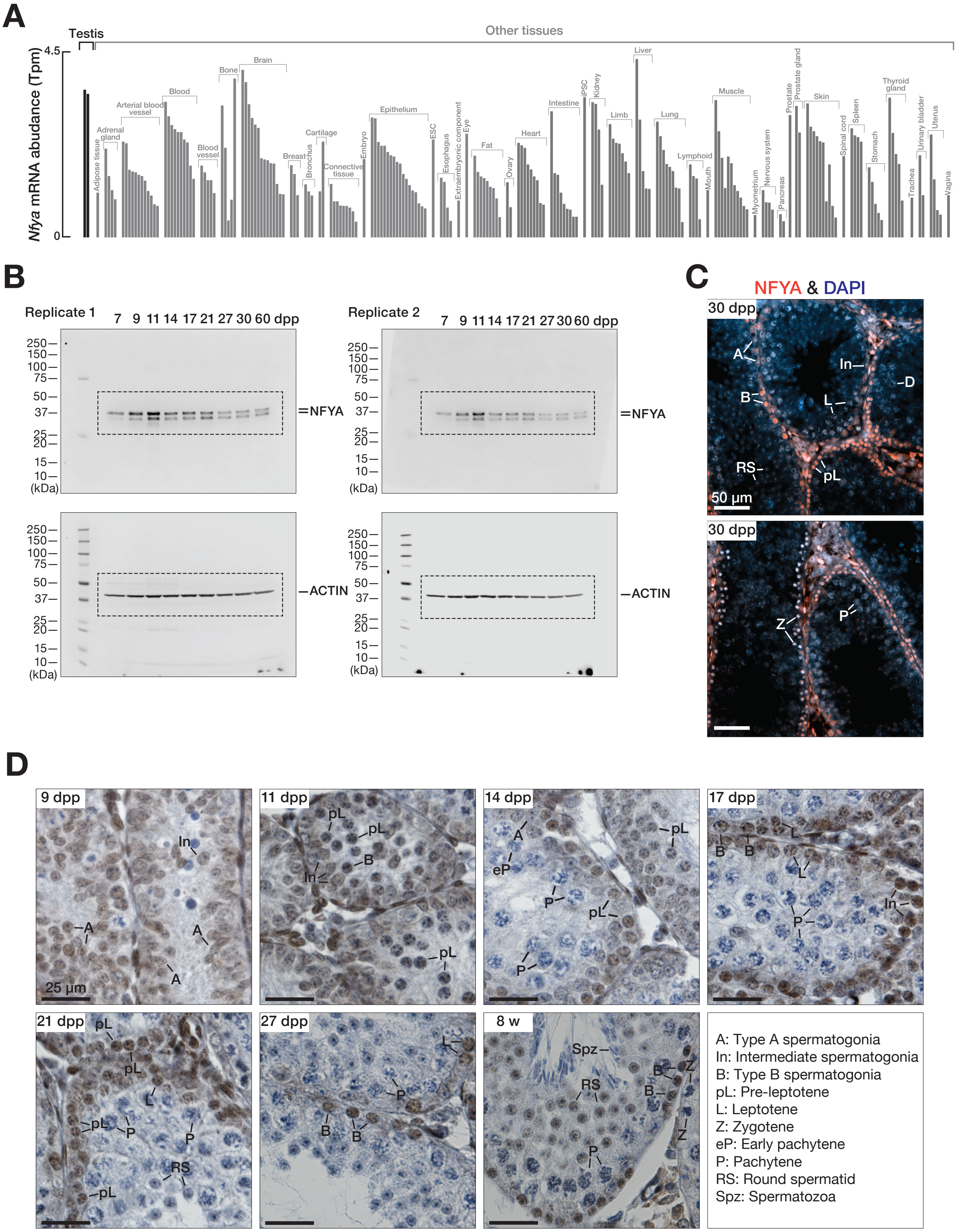

### Figure S8

Säflund et al., Figure S8

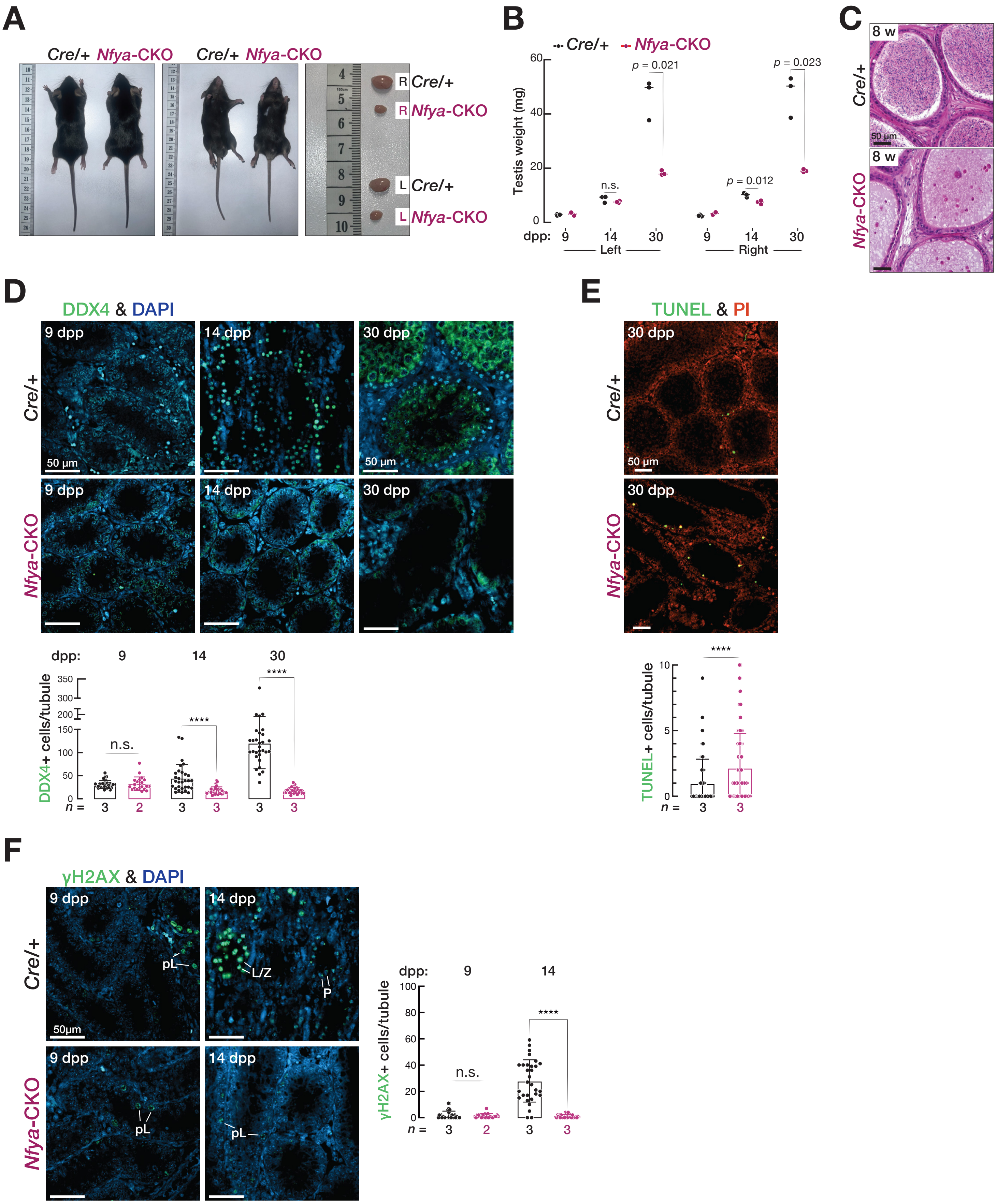

### Figure S9

Säflund et al., Figure S9

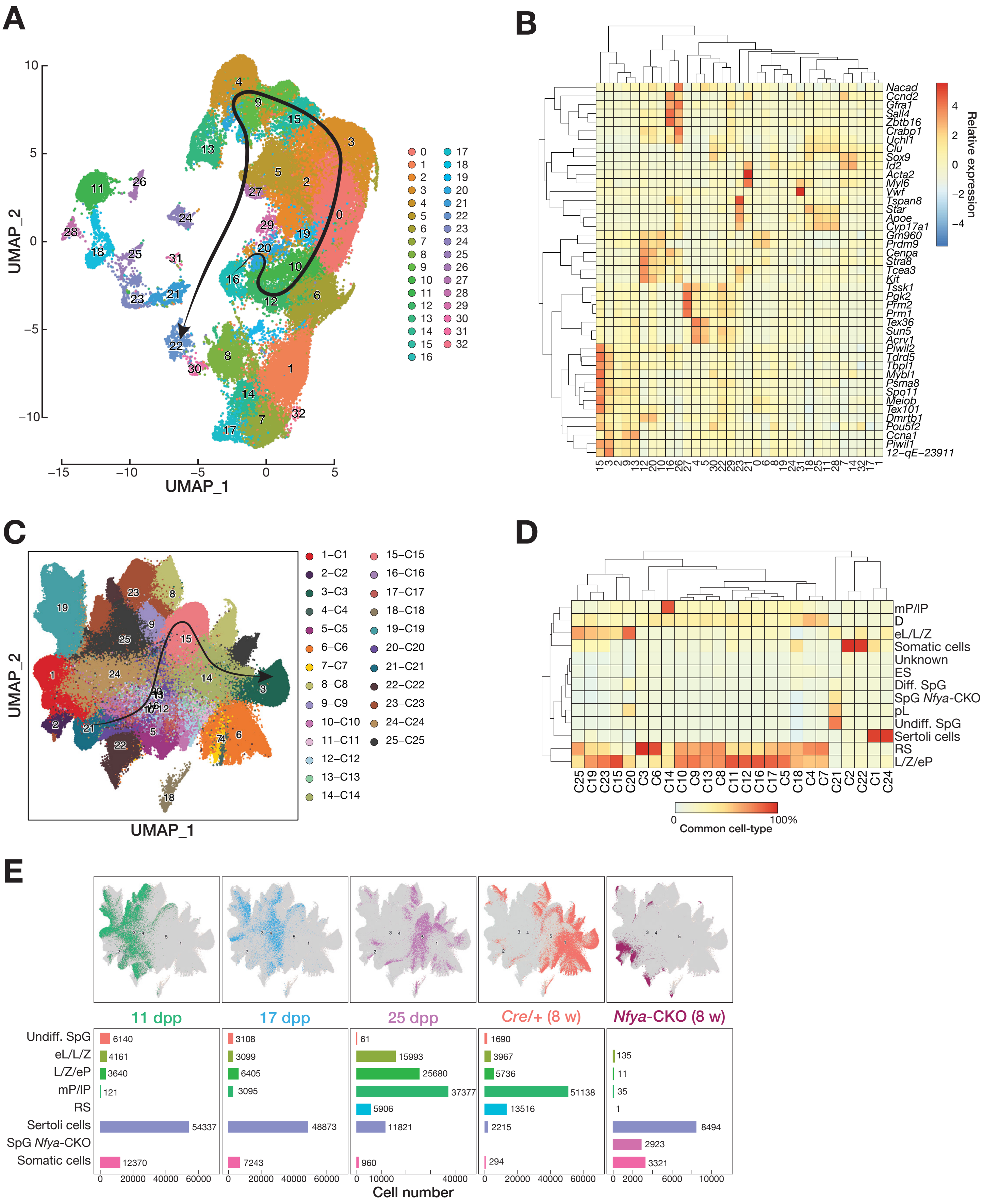
