## Supplementary material for "Transcription factor NFYA directs male meiotic entry by facilitating accessible chromatin at meiotic promoters in mice": Figure S3

Säflund et al., Figure S3

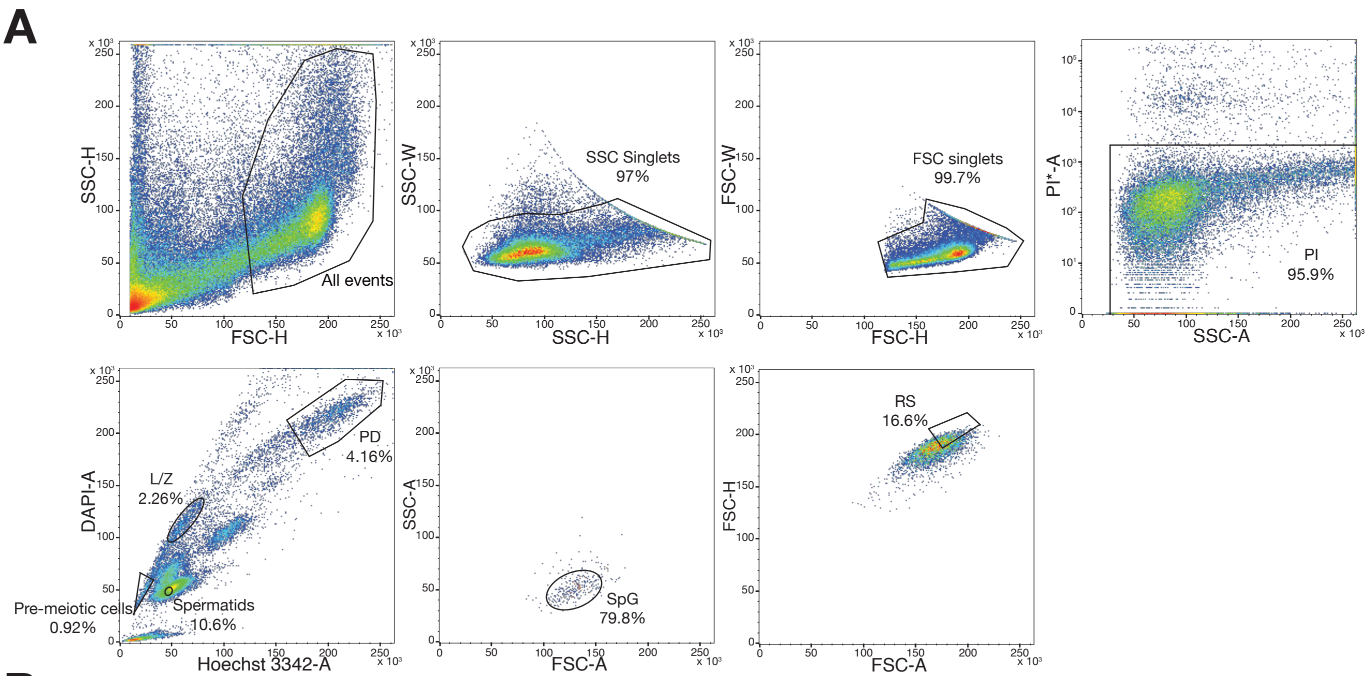


**B**

### SpG gate

| Animal # | GCNA & DMRT1 staining: GCNA$^+$/DMRT1$^+$ | GCNA & DMRT1 staining: GCNA$^-$/DMRT1$^-$ |
|---|---|---|
| 1 | 41 | 1 |
| 2 | 56 | 5 |
| 3 | 64 | 2 |
| 4 | 61 | 1 |
| 5 | 70 | 8 |
| 6 | 46 | 1 |
| Percent of each cell type | 95.3 ± 3.4 | 4.6 ± 3.4 |

Number of cells

### L/Z gate

SYCP3 & γH2A.X staining

| Animal # | SpG (SYCP3$^-$/γH2A.X$^-$) | L (SYCP3$^+$/γH2A.X$^+$) | Z (SYCP3$^+$/γH2A.X$^+$) | eP (SYCP3$^+$/γH2A.X$^+$) | P (SYCP3$^+$/γH2A.X$^+$) | D (SYCP3$^+$/γH2A.X$^+$) | RS (SYCP3$^-$/γH2A.X$^-$) | Sperm (SYCP3$^-$/γH2A.X$^-$) |
|---|---|---|---|---|---|---|---|---|
| 1 | 4 | 28 | 29 | 7 | 0 | 0 | 0 | 4 |
| 2 | 10 | 50 | 27 | 11 | 0 | 0 | 0 | 3 |
| 3 | 3 | 47 | 26 | 8 | 0 | 0 | 0 | 7 |
| 4 | 11 | 90 | 62 | 14 | 2 | 0 | 0 | 5 |
| 5 | 4 | 46 | 18 | 14 | 0 | 0 | 1 | 10 |
| 6 | 2 | 48 | 23 | 6 | 0 | 0 | 1 | 2 |
| Percent of each cell type | 5.2 ± 2.4 | 49.5 ± 5.8 | 29.4 ± 6.4 | 9.9 ± 2.6 | 0.2 ± 0.4 | 0 ± 0 | 0.4 ± 0.5 | 5.4 ± 3 |

Number of cells

### P/D gate

SYCP3 & γH2A.X staining

| Animal # | SpG (SYCP3$^-$/γH2A.X$^-$) | L (SYCP3$^+$/γH2A.X$^+$) | Z (SYCP3$^+$/γH2A.X$^+$) | eP (SYCP3$^+$/γH2A.X$^+$) | P (SYCP3$^+$/γH2A.X$^+$) | D (SYCP3$^+$/γH2A.X$^+$) | RS (SYCP3$^-$/γH2A.X$^-$) | Sperm (SYCP3$^-$/γH2A.X$^-$) |
|---|---|---|---|---|---|---|---|---|
| 1 | 1 | 0 | 0 | 0 | 27 | 21 | 0 | 0 |
| 2 | 0 | 13 | 0 | 0 | 41 | 32 | 1 | 0 |
| 3 | 0 | 0 | 0 | 0 | 30 | 35 | 0 | 0 |
| 4 | 0 | 0 | 0 | 0 | 47 | 35 | 21 | 0 |
| 5 | 3 | 5 | 0 | 0 | 61 | 46 | 12 | 4 |
| 6 | 0 | 3 | 1 | 2 | 51 | 50 | 8 | 0 |
| Percent of each cell type | 0.7 ± 1 | 3.6 ± 5.3 | 0.1 ± 0.3 | 0.3 ± 0.6 | 47.5 ± 3.5 | 41 ± 6.8 | 6.3 ± 7.2 | 0.5 ± 1.1 |

Number of cells

### RS gate

SYCP3 & γH2A.X staining

| Animal # | SpG (SYCP3$^-$/γH2A.X$^-$) | L (SYCP3$^+$/γH2A.X$^+$) | Z (SYCP3$^+$/γH2A.X$^+$) | eP (SYCP3$^+$/γH2A.X$^+$) | P (SYCP3$^+$/γH2A.X$^+$) | D (SYCP3$^+$/γH2A.X$^+$) | RS (SYCP3$^-$/γH2A.X$^-$) | Sperm (SYCP3$^-$/γH2A.X$^-$) |
|---|---|---|---|---|---|---|---|---|
| 1 | 0 | 0 | 0 | 0 | 0 | 0 | 80 | 0 |
| 2 | 0 | 0 | 0 | 0 | 0 | 0 | 35 | 14 |
| 3 | 0 | 0 | 0 | 0 | 0 | 0 | 71 | 13 |
| 4 | 0 | 0 | 0 | 0 | 0 | 0 | 48 | 14 |
| 5 | 0 | 0 | 0 | 0 | 0 | 0 | 83 | 3 |
| 6 | 0 | 0 | 0 | 0 | 0 | 0 | 95 | 0 |
| Percent of each cell type | 0 ± 0 | 0 ± 0 | 0 ± 0 | 0 ± 0 | 0 ± 0 | 0 ± 0 | 88.3 ± 11.2 | 11.6 ± 11.3 |

Number of cells

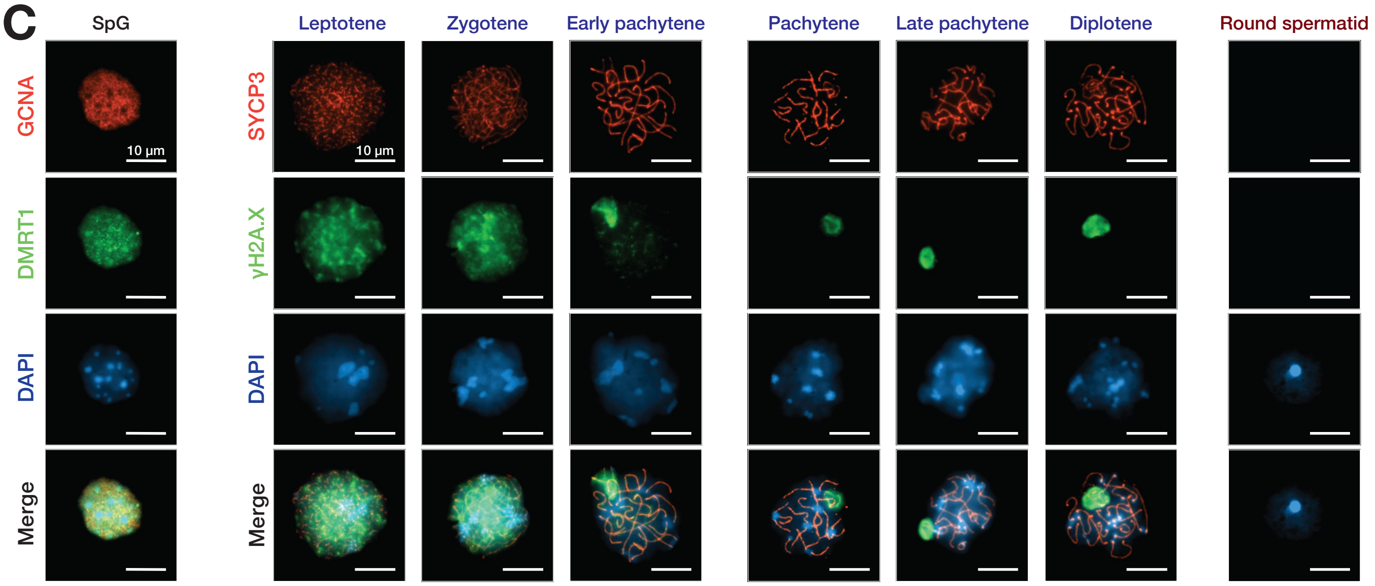
