## Supplementary material for "Transcription factor NFYA directs male meiotic entry by facilitating accessible chromatin at meiotic promoters in mice": Figure S6

Säflund et al., Figure S6

**A**

| Sample | NFYA CUT&RUN from SpG cells | | | | | NFYA CUT&RUN from L/Z cells | | | | | NFYA CUT&RUN from P/D cells | | | | |
|---|---|---|---|---|---|---|---|---|---|---|---|---|---|---|---|
| | # of NFYA peaks | % of peaks near TSSs | # of protein-coding genes | # of non-coding genes | % of peaks at enhancers (marked by H3K4me1) | # of NFYA peaks | % of peaks near TSSs | # of protein-coding genes | # of non-coding genes | % of peaks at enhancers (marked by H3K27Ac) | # of NFYA peaks | % of peaks near TSSs | # of protein-coding genes | # of non-coding genes | % of peaks at enhancers (marked by H3K27Ac) |
| Replicate 1 | 61,293 | 42.2% | 10,603 | 2,999 | 3.6% | 3,560 | 90.6% | 2,758 | 225 | 13.9% | 9,896 | 40.4% | 2,814 | 562 | 4.6% |
| Replicate 2 | 5,213 | 88.1% | 3,744 | 366 | 1.6% | 1,696 | 92.9% | 1,366 | 105 | 16.3% | 4,354 | 58.6% | 1,953 | 273 | 7.3% |
| Replicate 3 | 4,077 | 80.0% | 3,006 | 257 | 1.4% | 933 | 95.5% | 795 | 51 | 13.5% | 13,853 | 26.1% | 2,139 | 743 | 2.8% |

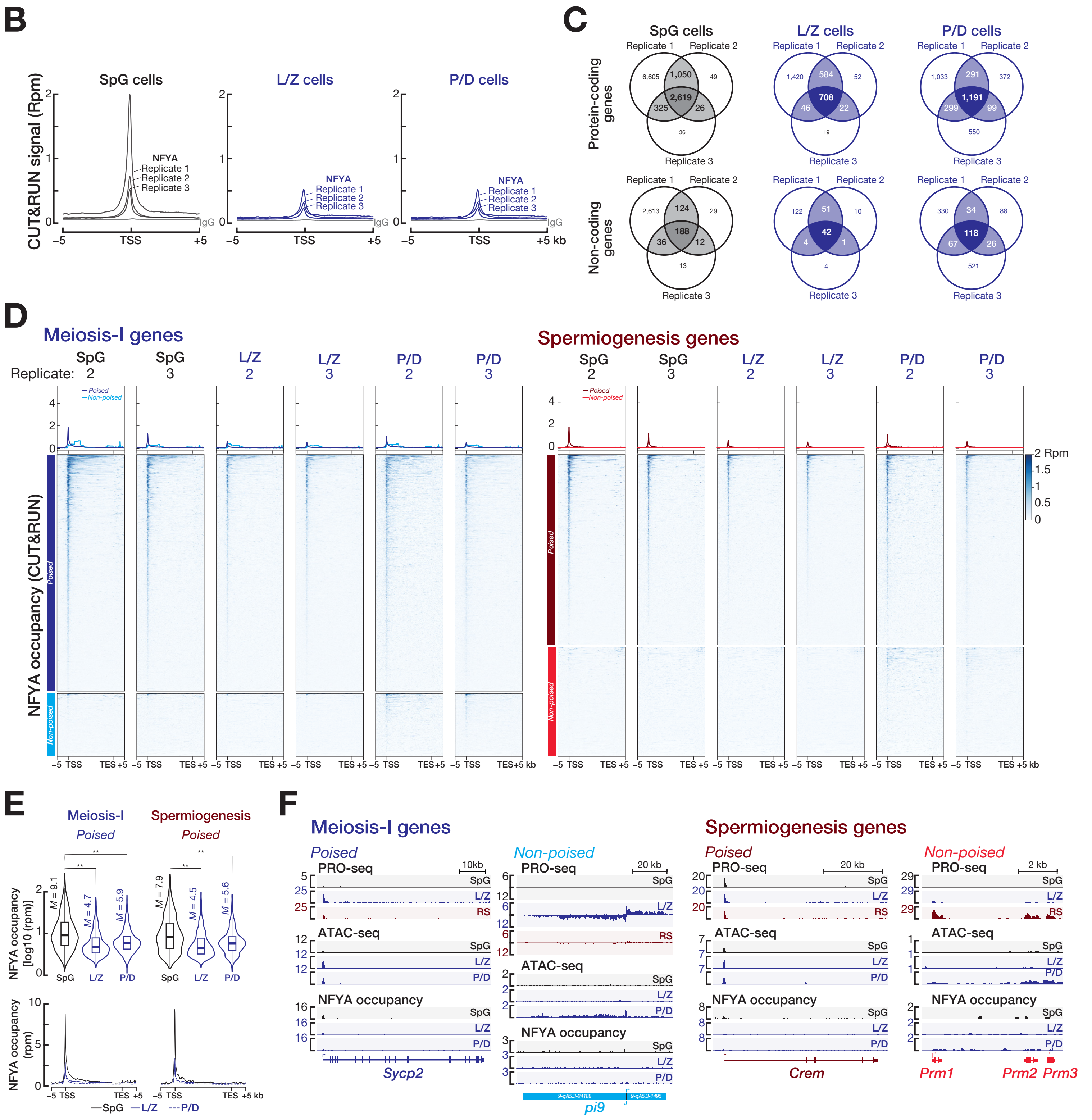
