## Supplementary material for "Transcription factor NFYA directs male meiotic entry by facilitating accessible chromatin at meiotic promoters in mice": Figure S7

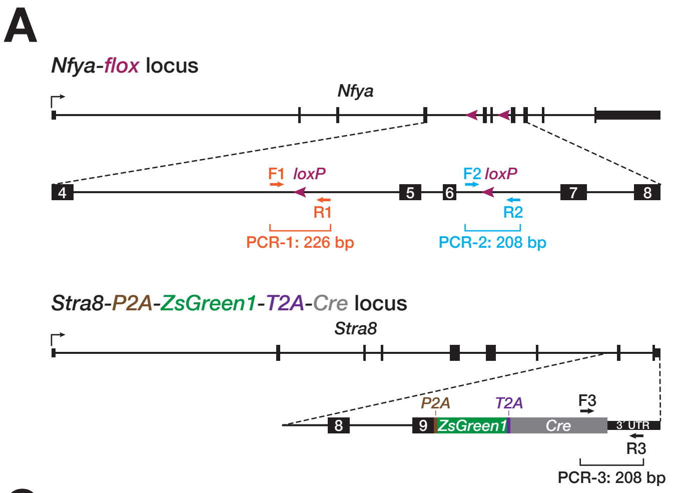


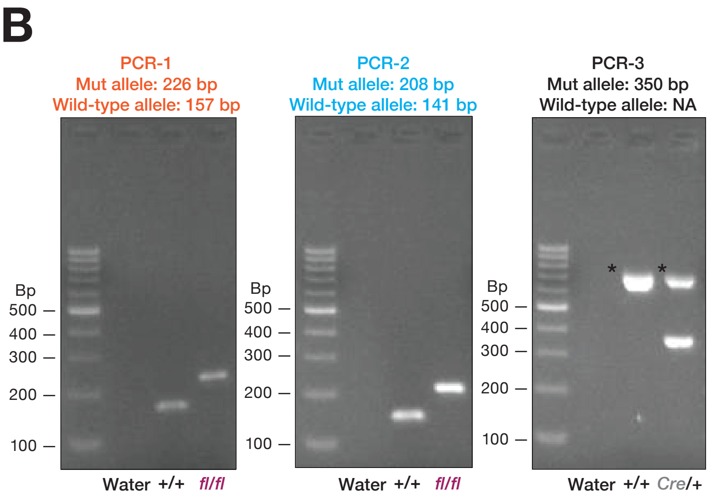


C

| Breeders #1 | ♂ Nfya$^{fl/+}$; Stra8-P2A-ZsGreen1-T2A-Cre$^{Cre/+}$ × ♀ Nfya$^{fl/+}$ and ♀ Nfya$^{fl/+}$ | | | | | |
|---|---|---|---|---|---|---|
| 122 total mice | fl/fl; +/+ | fl/fl; Cre/+ | fl/+; +/+ | fl/+; Cre/+ | +/+; +/+ | +/+; Cre/+ |
| Number of mice | 0 | 8 | 19 | 14 | 69 | 12 |
| Expected ratio | 0.125 | 0.125 | 0.25 | 0.25 | 0.125 | 0.125 |
| Observed ratio | 0 | 0.07 | 0.16 | 0.11 | 0.6 | 0.1 |

| Breeders #2 | ♂ Nfya$^{fl/+}$; Stra8-P2A-ZsGreen1-T2A-Cre$^{Cre/+}$ × ♀ Nfya$^{fl/+}$ and ♀ Nfya$^{fl/fl}$ | | | | | |
|---|---|---|---|---|---|---|
| 38 total mice | fl/fl; +/+ | fl/fl; Cre/+ | fl/+; +/+ | fl/+; Cre/+ | +/+; +/+ | +/+; Cre/+ |
| Number of mice | 9 | 3 | 8 | 12 | 1 | 5 |
| Expected ratio | 0.125/ 0.25 | 0.125/ 0.25 | 0.25 | 0.25 | 0.125/ 0 | 0.125/ 0 |
| Observed ratio | 0.24 | 0.08 | 0.21 | 0.32 | 0.03 | 0.13 |

| Breeders #3 | ♂ Nfya$^{fl/+}$; Stra8-P2A-ZsGreen1-T2A-Cre$^{Cre/+}$ × ♀ Nfya$^{fl/fl}$ and ♀ Nfya$^{fl/fl}$ | | | | | |
|---|---|---|---|---|---|---|
| 22 total mice | fl/fl; +/+ | fl/fl; Cre/+ | fl/+; +/+ | fl/+; Cre/+ | +/+; +/+ | +/+; Cre/+ |
| Number of mice | 2 | 6 | 7 | 7 | 0 | 0 |
| Expected ratio | 0.25 | 0.25 | 0.25 | 0.25 | 0 | 0 |
| Observed ratio | 0.09 | 0.27 | 0.32 | 0.32 | 0 | 0 |

| Breeders #4 | ♂ Nfya$^{fl/fl}$ × ♀ Nfya$^{fl/+}$; Stra8-P2A-ZsGreen1-T2A-Cre$^{Cre/+}$ and ♀ Nfya$^{fl/+}$; Stra8-P2A-ZsGreen1-T2A-Cre$^{Cre/+}$ | | | | | |
|---|---|---|---|---|---|---|
| 22 total mice | fl/fl; +/+ | fl/fl; Cre/+ | fl/+; +/+ | fl/+; Cre/+ | +/+; +/+ | +/+; Cre/+ |
| Number of mice | 9 | 5 | 5 | 3 | 0 | 0 |
| Expected ratio | 0.25 | 0.25 | 0.25 | 0.25 | 0 | 0 |
| Observed ratio | 0 | 0.07 | 0.16 | 0.11 | 0.6 | 0.1 |

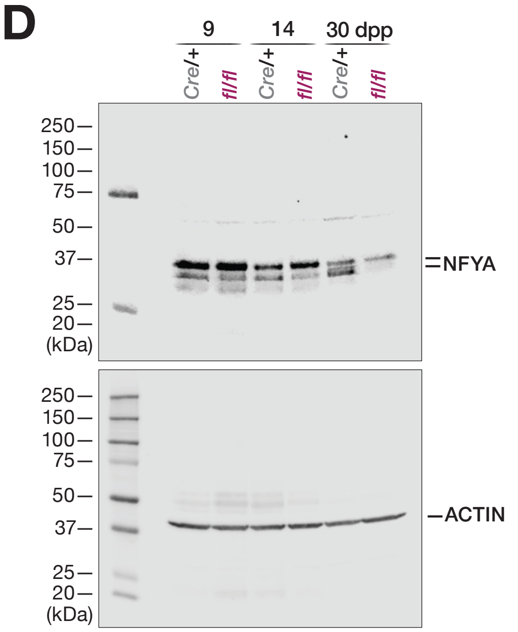


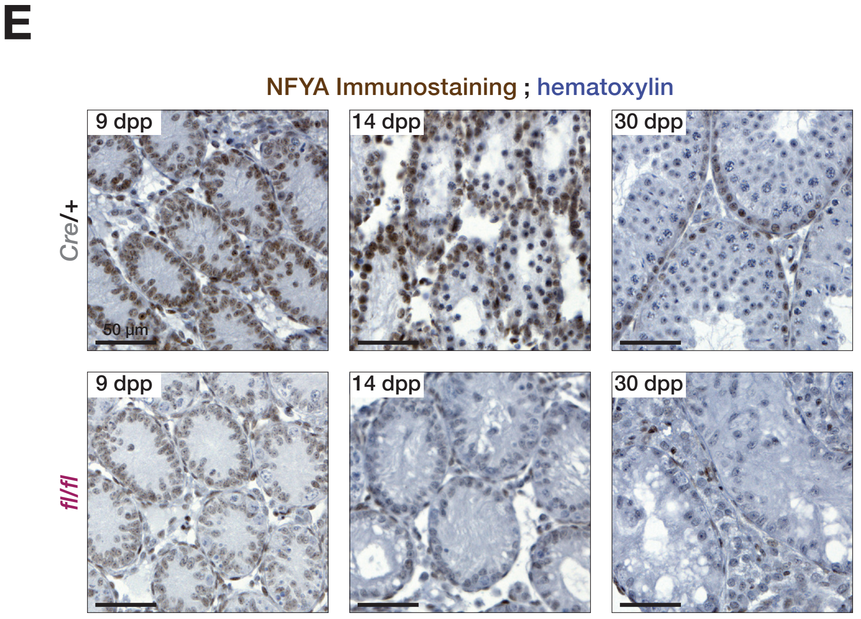
