## Supplementary material for "Transcription factor NFYA directs male meiotic entry by facilitating accessible chromatin at meiotic promoters in mice": Legends for supplemental figures

Supplementary Materials for

**Transcription factor NFYA directs male meiotic entry by facilitating accessible chromatin at the promoters of genes expressed during meiosis**

Martin Säflund, Masomeh Askari, Atiyeh Eghbali, Mukhtar Mohamed Abdi, Ann-Kristin Östlund Farrants, Tianxiong Yu,** Deniz M. Özata*

Supplemental Figure S1 related to Figure 1.

Each major step of spermatogenesis retains a distinct gene expression profile.

**(A)** Heatmaps show absolute steady-state transcript abundance of mitosis, meiosis-I and spermiogenesis genes in spermatogonia (SpG), pachytene and diplotene (P/D), secondary spermatocytes (SpII) and round spermatids (RS). Genes whose RNA transcript abundance is ≥4-fold higher in SpG compared to P/D, SpII and RS are classified as mitosis specific genes. Meiosis-I genes are classified as those whose RNA transcript abundance is ≤0.25-fold lower in SpG compared to P/D and are ≥4-fold higher in P/D compared to SpII. Spermiogenesis genes are defined as those with ≤2-fold increases in SpII and RS compared to P/D and ≤0.25-fold lower in SpG compared to P/D.

**(B- D)** Integrated Genomics Viewer (IGV) views of reads per kilobase million (Rpkm)-normalized RNA sequencing signal for gene examples from **(B)** Mitosis genes, *Sall4, Dmrt1*, **(C)** Meiosis-I genes, *pi9*, *Sycp2*, **(D)** spermiogenesis genes, *Crem*, *Spata6*.

**(E)** Gene Ontology (GO) analysis of meiosis-I (top) and spermiogenesis (bottom) genes. The gene categories are sorted by fold enrichment.

Supplemental Figure S2 related to Figure 1.

**Paused polymerase accumulates around the promoters of genes expressed during meiosis in spermatogonia.**

**(A)** Violin plots represent the distance (bp) from TSS to nearest promoter PRO-seq signal of poised and non-poised meiosis-I (left) and spermiogenesis (right) genes in spermatogonia (SpG), primary spermatocytes (SpI) and round spermatids (RS). Horizontal lines represent mean. Whiskers represent maximum and minimum values. Interquartile range (IQR) represented by boxplots.

**(B)** Heatmaps show the relative promoter (left; from TSS to first 5% of gene length) and gene body (right; from first 30% of gene length to TES) PRO-seq signal for mitosis genes in spermatogonia (SpG), primary spermatocytes (SpI) and round spermatids (RS).

**(C)** Line plots represent the average promoter (dash line with triangle) and gene body (line with squares) relative PRO-seq signal for mitosis genes across SpG, SpI and RS cells.

**(D)** Violin plots show reads per million (Rpm)-normalized A-MYB (left) and BRDT (right) occupancies around the promoters of mitosis, poised and non-poised meiosis-I and spermiogenesis genes in pachytene spermatocytes. Horizontal lines represent median. Whiskers represent maximum and minimum values. IQR represented by boxplots.; ****p < 0.0001; Mann-Whitney-Wilcoxon U test.

**Supplemental Figure S3 related to Figures 2 and 3.**

**Strategy for purifying male germ cells using fluorescence-activated cell sorter (FACS).**

**(A)** Gating strategy used to purify spermatogonia (SpG), leptotene/zygotene spermatocytes (L/Z), pachytene/diplotene spermatocytes (P/D) and round spermatids (RS). Cells were stained with Hoechst 3342 and propidium iodide (PI). 488-nm laser was used to excite PI, while 405-nm laser was used to excite Hoechst 33342. The emission for PI was recorded using 700/54 nm bandpass filter, while the emission of Hoechst 33342 was recorded by 660/10-nm (red) and 448/45-nm (blue) bandpass filters.

**(B)** Tables summarizes counted cells per sorted gate. Six independent replicates were assessed per gate and the average percentage of each cell type calculated with standard deviation.

**(C)** Immunofluorescence staining for various germ cell markers from the chromosome spreads of four FACS-purified populations (SPG, L/Z, P/D, RS) to assess the purify of each gate. Anti-GCNA and anti-DMRT1 to assess SpG; anti-SYCP3, anti-γH2A.X to assess L/Z, P/D and RS gates. Cell pictures are ordered from left to right according to the trajectory of male germ cell development. Scale bars: 10 µm.

**Supplemental Figure S4 related to Figures 2.**

**Poised genes have accessible chromatin around their promoter regions in spermatogonia.**

**(A)** Metagene plots (top) and heatmaps (bottom) show reads per million (Rpm)-normalized Assay for Transposase-Accessible Chromatin sequencing (ATAC-seq) signals in the –5 kb to +5 kb window flanking transcription start sites (TSSs) and transcription end sites (TESs) of poised and non-poised genes from the second and third replicates of spermatogonia (SpG), leptotene/zygotene (L/Z), and pachytene/diplotene (P/D). Embryonic stem cells (ESC^40^) served as control.

**(B)** Heatmap shows spearman’s correlation between the biological replicates of ATAC-seq from each cell population. Agreement between replicates; SpG Spearman’s ρ > 0.83, L/Z Spearman’s ρ > 0.92, P/D Spearman’s ρ > 0.95).

**Supplemental Figure S5 related to Figure 3.**

**Temporal expression of NFYA in the testes of wild-type mouse.**

**(A)** Bars shows transcripts per million (Tpm)-normalized mRNA abundance of *Nfya* across different mouse tissues. Data is from mouse Encyclopedia of DNA Elements (ENCODE).

**(B)** Western blot shows NFYA protein abundance in the testes from stage mice (i.e., 7, 9, 11, 14, 17, 21, 27, 30 and 60 days postpartum [dpp]). ACTIN used as loading control. Each lane contained 75 µg total testis protein isolate.

**(C)** Immunofluorescence staining against NFYA on the testis section from 30 dpp mouse. A, type A spermatogonia; In, intermediate spermatogonia; B, type B spermatogonia; pL, preleptotene; L, leptotene; Z, zygotene; P, pachytene; RS, round spermatid. Scale bars: 50 µm.

**(D)** Immunohistochemical (IHC) detection of NFYA on Bouin fixed testes sections from stage mice. A, type A spermatogonia; In, intermediate spermatogonia; B, type B spermatogonia; pL, preleptotene; L, leptotene; Z, zygotene; eP, early pachytene; P, pachytene; RS, round spermatid; Spz, Spermatozoa.

**Supplemental Figure S6 related to Figure 3.**

**NFYA binds the promoters of poised genes in spermatogonia.**

**(A)** Tables summarize genome-wide significant NFYA CUT&RUN peaks in the three replicates of spermatogonia (SpG), leptotene/zygotene spermatocytes (L/Z), and pachytene/diplotene spermatocytes (P/D).

**(B)** Metaplots show average reads per million (Rpm)-normalized NFYA CUT&RUN signals in the –5 kb to +5 kb window flanking transcription start sites (TSSs) of all mouse RefSeq genes in SpG, L/Z, and P/D. IgG used for negative control.

**(C)** Venn diagrams show number of overlapping protein- and non-coding genes across three biological replicates.

**(D)** Metagene plots (top) and heatmaps (bottom) show Rpm-normalized NFYA CUT&RUN signals in the –5 kb to +5 kb window flanking transcription start sites (TSSs) and transcription end sites (TESs) of poised and non-poised genes in the second and third replicates from SpG, L/Z, P/D.

**(E)** Violin plot (top; on log10 scale) and metaplot (bottom) show NFYA occupancy around the promoters of poised meiosis-I (left) and spermiogenesis (right) genes in SpG, L/Z, and P/D. Horizontal lines represent median. Whiskers represent maximum and minimum values. IQR represented by boxplots. ***p* < 0.01; two-sided wilcoxon matched-pairs signed ranked sum test.

**(F)** IGV tracks of PRO-seq signal, ATAC-seq signal, and NFYA occupancy (CUT&RUN) at gene boundaries of exemplified poised and non-poised meiosis-I (*Sycp2, pi9*) and spermiogenesis (*Crem, Prm1, Prm2, Prm3*) genes from SpG, L/Z, P/D, and RS.

**Supplemental Figure S7 related to Figure 4.**

**Male germ cell-specific deletion of *Nfya*.**

**(A)** Strategy to delete *Nfya* in male germ cells using Stra8-Cre-loxP system.

**(B)** Agarose gel electrophoresis images for genotyping of mutants by PCR. Each *loxp* site flanking the exon was validated by PCR amplification (PCR-1 and PCR-2 primers are highlighted in red and light blue) in mice with *Stra8-Cre* transgene which was also confirmed by PCR (PCR-3 primers are highlighted in black). * Represents internal control PCR bands.

**(C)** Tables show number and frequency of pups with all possible genotypes from four different breading pairs. Highlighted are the number of pups with homozygous deletion.

**(D)** Western blot images show NFYA abundance in *Cre/*+ and *fl/fl* mice at 9, 14 and 30 days postpartum (dpp)*.* ACTIN served as loading control. Each lane contained 75 µg testis protein.

**(E)** Immunohistochemical (IHC) detection of NFYA on Bouin fixed testes sections from from *Cre/*+ and *fl/fl* mice at 9, 14, 30 dpp. Scale bars: 50 µm.

**Supplemental Figure S8 related to Figure 4.**

**NFYA deficient male germ cells arrest at meiotic entry.**

**(A)** Comparison of testes from 8-week-old *Cre/+* and *Nfya-CKO* mice.

**(B)** Relative testes weights in *Cre/+* and *Nfya-CKO* mice at 9 days postpartum (dpp), 14 dpp, and 30 dpp. (*n* = 2 or *n* = 3). Significance is measured using two-sided paired student t-test.

**(C)** Hematoxylin and eosin staining of the caudal epididymis sections from 8-week-old *Cre/+* and *Nfya-CKO* mice. Scale bars: 50 μm.

**(D-F)** Immunofluorescence staining of testes sections from 9, 14 and 30 dpp *Cre/*+ and *Nfya*-CKO for **(D)** DDX4, **(E)** TdT-mediated dUTP Nick-End Labeling (TUNEL) and **(F)** γH2A.X. Bars represent quantification for positive cells per tubule from three independent biological replicates. Whiskers represent standard deviation. Significance is measured using two-sided unpaired student t-test. Scale bars: 50 μm.

**Supplemental Figure S9 related to Figures 5, 6, and 7.**

**Identification of cell populations of the testes from 8-week-old Cre/+ and Nfya-CKO mice along with wild-type staged mice using simultaneous single-cell (sc) RNA-seq and ATAC-seq.**

**(A)** Uniform manifold approximation and projection (UMAP) space visualizes 32 clusters obtained by dimension reduction analysis on 45,250 individual cells from 11, 17, 25 dpp, *Cre/+* and *Nfya-CKO* mice. Developmental trajectory for germ cells is shown.

**(B)** Unsupervised hierarchical clustering of 32 clusters based on the relative expression of cell markers.

**(C)** UMAP space visualizes 25 cell clusters defined by the chromatin accessibility obtained from scATAC-seq of 343,866 individual cells from 11, 17, 25 dpp, *Cre/+* and *Nfya-CKO* mice. Developmental trajectory for germ cells is shown.

**(D)** Heatmap shows the percentage of cell, whose type is defined by scRNA-seq, in clusters obtained by scATAC-seq.

**(E)** UMAP spaces (top) illustrate identity of cellular diversity obtained from scATAC-seq in each mouse compared to the background of remaining mice (grey color). Bars (bottom) represent the total number of individual cells belonging to each major cell population.

Supplemental Table S1

**Gene category lists and PRO-seq peak distance to TSS of genes from the gene categories.**

**(A)** Absolute transcript abundance of all genes in FACS-sorted C57BBL/6 mice germ cells. Data from Cecchini et al., Reproduction, 2023.

**(B)** Mean distance from TSS to summit of nearest PRO-seq peak for Meiosis-I and Spermigoenesis genes.

**(C, D, E)** Gene ontology analysis for genes classified as mitosis, meiosis I and spermiogenesis genes.

**Supplemental Table S2.**

**Examination of accessible chromatin around the promoters of poised and non-poised genes in spermatogonia, leptotene/zygotene and pachytene/diplotene cells. Mitosis specific genes are listed and genes used as negative control.**

**Supplemental Table S3.**

**Distance from nearest NFYA peak (CUT&RUN) to the promoter of mitosis, meiosis I and spermiogenesis genes. In spermatogonia, leptotene/zygotene and pachyeten/diplotene cells.**

**Supplemental Table S4.**

**Number of reads and list of genes captured during scRNA-seq and scATAC-seq.**

**(A)** Type of cells, and their numbers as well as number of genes captured by scRNA-seq.

**(B)** Expression of genes in the different cell types captured by scRNA-seq.

**(C)** Captured reads from scATAC-seq libraries.

**(D)** Average scATAC-seq signal in undifferentiated spermatogonia from *Cre/+* and *Nfya*-CKO mice.

**Supplemental Table S5.**

**Statistics for sequenced libraries used in this study.**
